# Major alleles of *CDCA7α* shape CG-methylation in *Arabidopsis thaliana*

**DOI:** 10.1101/2025.09.03.673934

**Authors:** Pierre Bourguet, Zdravko J Lorković, Darya Kripkiy Casado, Valentin Bapteste, Chung Hyun Cho, Anna Igolkina, Cheng-Ruei Lee, Magnus Nordborg, Frédéric Berger, Eriko Sasaki

## Abstract

DNA methylation is a key epigenetic mark that impacts gene expression and represses transposable elements (TEs) in eukaryotes. Numerous examples of *cis*-elements targeted by DNA methylation, particularly at CG sites (mCG), have been reported to be under selective pressure in animals and plants. By contrast, there is limited knowledge of *trans*-regulators of mCG leading to adaptation. Here, using genome-wide association studies, we identify CELL DIVISION CYCLE-ASSOCIATED PROTEIN 7 ALPHA (*CDCA7α*) as a *trans*-regulator of mCG in natural populations of *Arabidopsis thaliana*. CDCA7α and its paralog, CDCA7β, directly bind to the chromatin remodeler DECREASE IN DNA METHYLATION 1 (DDM1), which facilitates access of methyltransferases to DNA. CDCA7α/β selectively regulates mCG and minimally impacts other DDM1-dependent processes such as non-CG methylation and histone variant deposition. We identify the *cis*-regulatory sequence modulating *CDCA7α* expression in natural populations and determining the degree of mCG and TE silencing. The geographic distribution of *CDCA7α* alleles suggests that new alleles have repeatedly expanded to novel ecological niches, indicating a potential role in local adaptation. Altogether, our findings provide new insight into how changes in global DNA methylation levels through transcriptional regulation of the epigenetic machinery have the capacity to facilitate local adaptation.

## Introduction

5-methylcytosine (5mC) is the predominant form of DNA modification in multicellular eukaryotes (Boulias and Greer, 2022; Romero Charria *et al*., 2024). 5mC regulates gene expression during development and preserves genome integrity by repressing transposable elements (TEs) (Smith and Meissner, 2013; Schmitz, Lewis and Goll, 2019). While conserved molecular pathways control the genomic distribution of 5mC (Law and Jacobsen, 2010; Greenberg and Bourc’his, 2019), the epigenome varies between individuals due to genetic differences, environmental influences and stochastic catalytic activity (Kader and Ghai 2017; Fraga et al. 2005; Jeltsch and Jurkowska 2014; Johannes and Schmitz 2019; Briffa et al. 2023; Richards 2006). In many organisms, epigenome variation contributes to phenotypic variation at the cellular, individual and population level (Becker *et al*., 2011; Chatterjee *et al*., 2015; Quadrana and Colot, 2016; Zheng *et al*., 2017; Schmid *et al*., 2018; Greenberg and Bourc’his, 2019; Venney *et al*., 2023). Key remaining questions are how this epigenome variation was established and whether it contributes to adaptive evolution in nature.

Proteins essential for 5mC maintenance are well characterized in plants (Zhang, Lang and Zhu, 2018). Methylation of CG dinucleotides (mCG) is maintained by METHYLTRANSFERASE 1 (MET1), while non-CG methylation (mCH, where H is A, T or C) requires the chromomethyltransferases (CMT) CMT2 and CMT3, and the RNA-directed DNA methylation (RdDM) pathway. MET1 and CMT2/3 rely on the chromatin remodeler DECREASE IN DNA METHYLATION 1 (DDM1), which facilitates methyltransferase access to DNA by remodeling nucleosomes.

In wild plant populations, 5mC levels are often associated with local climates (Foust et al. 2016; Platt et al. 2015; Sammarco et al. 2024; Dubin et al. 2015; Rodríguez et al. 2022). The 1001 Epigenomes Project revealed 5mC variation for 1107 worldwide accessions of *Arabidopsis thaliana*, and genome-wide association studies (GWAS) demonstrated that at least some of it had a genetic basis (Kawakatsu *et al*., 2016). mCH variation is controlled by genetic variation at *trans*-acting modifiers, including *CMT2*, *CMT3*, and *NRPE1*, a subunit of RNA polymerase V (Kawakatsu *et al*., 2016; Sasaki *et al*., 2019, 2022). These regulators influence TE activity and their allelic distribution shapes geographic clines (Baduel *et al*., 2021; Sasaki *et al*., 2022). *Trans* regulators of mCG variation in gene bodies have been studied (Zhang *et al*., 2020; Briffa *et al*., 2023; Pisupati *et al*., 2023), although the function of gene body methylation remains debated (Muyle *et al*., 2022). In contrast, the genetic basis of mCG variation at TEs remains unknown, despite its critical role in transcriptional silencing and genome stability.

Here, we investigated *trans* regulators of mCG variation at TEs in natural *A. thaliana* lines. Using GWAS, we identified *CDCA7α*, an ortholog of the human *Cell division cycle-associated protein 7* (Thijssen *et al*., 2015), as a major *trans* regulator of mCG variation. We show that CDCA7α and its paralog, CDCA7β, act as facultative activators of DDM1 and specifically promote its function in depositing mCG. In natural populations, the highly diverged *CDCA7α* promoter region modulates its expression, thus influencing the degree of mCG and TE silencing genome-wide. New *CDCA7α* alleles emerged multiple times after speciation and accompanied expansion to new ecological niches, suggesting a role of mCG optimization in local adaptation.

## Results

### Natural TE mCG is controlled by genetic variants of ***CDCA7α***

To explore the genetic architecture of mCG variation, we conducted GWAS using the 1001 Epigenomes data set (*n*=774) (Kawakatsu *et al*., 2016). We focused on mCG levels from two domains of chromatin in which TEs are repressed by distinct mechanisms: euchromatic regions, represented by RdDM-targeted TEs, and heterochromatic regions, represented by CMT2-targeted TEs (Kawakatsu et al. 2016; see also methods) (**Fig. 1a, S1a**).

**Figure 1.**
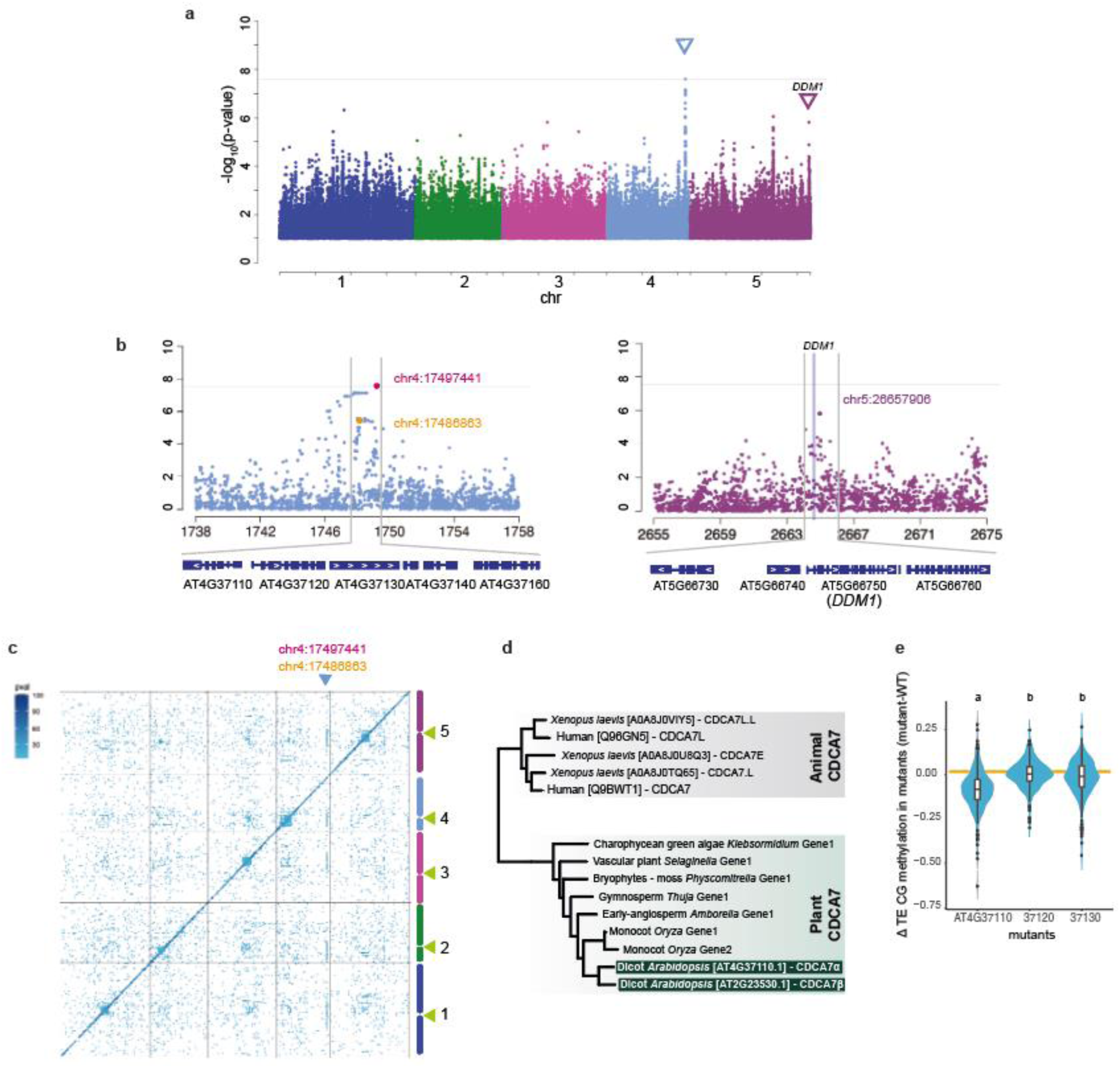
The genetic basis of CG methylation. **a,** GWAS for mCG on CMT2-targeted TEs. **b,** Zoom-in figures show the highest peak (left) and the *DDM1* region (right). **c,** Association for mCG on individual TEs. Associations with -log_10_*p*-value > 6 are shown. Green triangles indicate centromeric regions. **d,** Simplified phylogenetic tree of CDCA7 in vertebrates and plants. The full tree is shown in Fig. S3. **e,** Differential mCG levels of TEs associated with chr4:17486863 or chr4:17497441 for loss-of-function mutants. Lower-case letters indicate significant differences as determined using Tukey’s HSD test (*p* < 0.05).

While no genetic variant was associated with average mCG in RdDM-targeted TEs, several peaks were detected in GWAS for average mCG levels in CMT2-targeted TEs on chromosomes (chr) 4 and 5 (**Fig. 1a, S1a-e, g, Supplementary Table S1**). The association on the right end of chr 5 (chr5:26657906) was not statistically significant but was located just 4.8 kbp downstream of the coding region of *DDM1* and covered the coding to the 3’ region of *DDM1* (-log_10_*p*-value=5.80; Minor Allele Frequency (MAF)=47%) (**Fig. 1b, S1j**). This peak may reflect several causative alleles and one allele is associated with a SNP on chr5:26649074 resulting in an amino acid substitution (V9F) in the first exon of *DDM1*. This amino acid is not conserved across Angiosperms (**Fig. S2a**), does not belong to a domain of known function (**Fig. S2b**), and is not represented in available cryo-EM structures. Given the known major role of *DDM1* in the control of mCG at CMT2-dependent TEs (Zemach *et al*., 2013; Akinmusola, Wilkins and Doughty, 2023), a peak in *DDM1* is not unexpected, and the association is unlikely to be spurious.

The significant peak on chr4 spanned a 15 kbp region from chr4:17482000 to 17497000, and appeared to involve two distinct haplotypes (**Fig. 1b**). One haplotype was associated with the highest peak (chr4:17497441; -log_10_p-value=7.6; MAF=5.3%), and a complete subset of the other haplotype was associated with the other SNP chr4:17486863 (-log_10_p-value=5.47, MAF=30.1%). GWAS accounting for the genetic variants at chr4:17497441 did not affect the peak at *DDM1*; i.e., the associations were independent (**Fig. S1f, h, i**).

To gain further insight, we conducted GWAS for mCG levels of 6379 individual TEs commonly found in all 774 lines and containing at least one CG site (Sasaki *et al*., 2019). Strong associations were primarily observed in *cis* (diagonal), suggesting either direct epigenetic inheritance of mCG, or that genetic variation within TEs influences mCG levels, consistent with a previous study on differential mCG variation (Hüther *et al*., 2022). In addition, the vertical lines on the graph identified clear *trans* regulation of mCG levels in TEs distributed around pericentromeric regions associated with the chromosome 4 SNPs discussed above (**Fig. 1c**). Chr4:17486863 and chr4:17497441 showed associations with 229 and 116 TEs (threshold -log_10_*p*-value>6, MAF>5%), respectively, with a significant overlap (chr4:17486863: 62% and chr4:17497441: 75%) with CMT2-targeted TEs and almost no overlap (5.2% and 0%) with RdDM-targeted TEs.

Two haplotypes represented by chr4:17486863 and chr4:17497441 included several genes, but were centered on *AT4G37110* and *AT4G37120*. *AT4G37120* encodes SWELLMAP 2 (SMP2), a step II splicing factor that works redundantly with the homolog *SMP1* (Liu, Wu and Zhang, 2016). *AT4G37110* contains a zinc finger domain with homology to human Cell Division Cycle Associated 7 (CDCA7) (Thijssen *et al*., 2015), which has been shown to recruit the vertebrate ortholog of DDM1, “Helicase, Lymphoid-Specific” (HELLS; (Jenness *et al*., 2018; Unoki, 2021; Shinkai *et al*., 2024; Wassing *et al*., 2024), making it an excellent candidate for being the causal gene (**Fig. 1d**).

Further evidence for this hypothesis was provided by bisulfite sequencing of loss-of-function mutants for *SMP2*, *AT4G37110,* and the neighbouring gene *AT4G37130* as control (**Fig. 1e**). Unlike the other genes, the *at4g37110* mutant showed significantly lower mCG levels in the same TEs identified as targets of the *trans* modifier using GWAS (**Fig. 1e**). These results strongly suggest that allelic variation of *AT4G37110* is responsible for the identified association on chromosome 4 and that *AT4G37110* is an ortholog of human *CDCA7*.

### CDCA7α and CDCA7β redundantly maintain mCG and heterochromatin features to enforce silencing of TEs

In addition to *AT4G37110*, *AT2G23530* also contains a zinc finger domain with high sequence homology to metazoan *CDCA7*, suggesting that there might be two *bona fide CDCA7* orthologs in *A. thaliana* (Funabiki *et al*., 2023). Our study confirms this (see below). Phylogenetic analysis revealed that *CDCA7* duplication occurred in Brassicaceae (**Fig. S3a, b**), coinciding with a whole-genome duplication event (Schranz, Mohammadin and Edger, 2012). These two putative *A. thaliana CDCA7* orthologs share similar expression patterns throughout development (**Fig. S4**). Given its higher expression, *AT4G37110* was designated *CDCA7α*, while *AT2G23530* was named *CDCA7β* (**Fig. 2a**). According to ancestral sequence reconstruction, the zinc finger domain of *CDCA7β* is more similar to the ancestral gene than *CDCA7α* (**Fig. S3c, d**). We found no evidence for allelic variation affecting mCG in *CDCA7β*.

**Figure 2.**
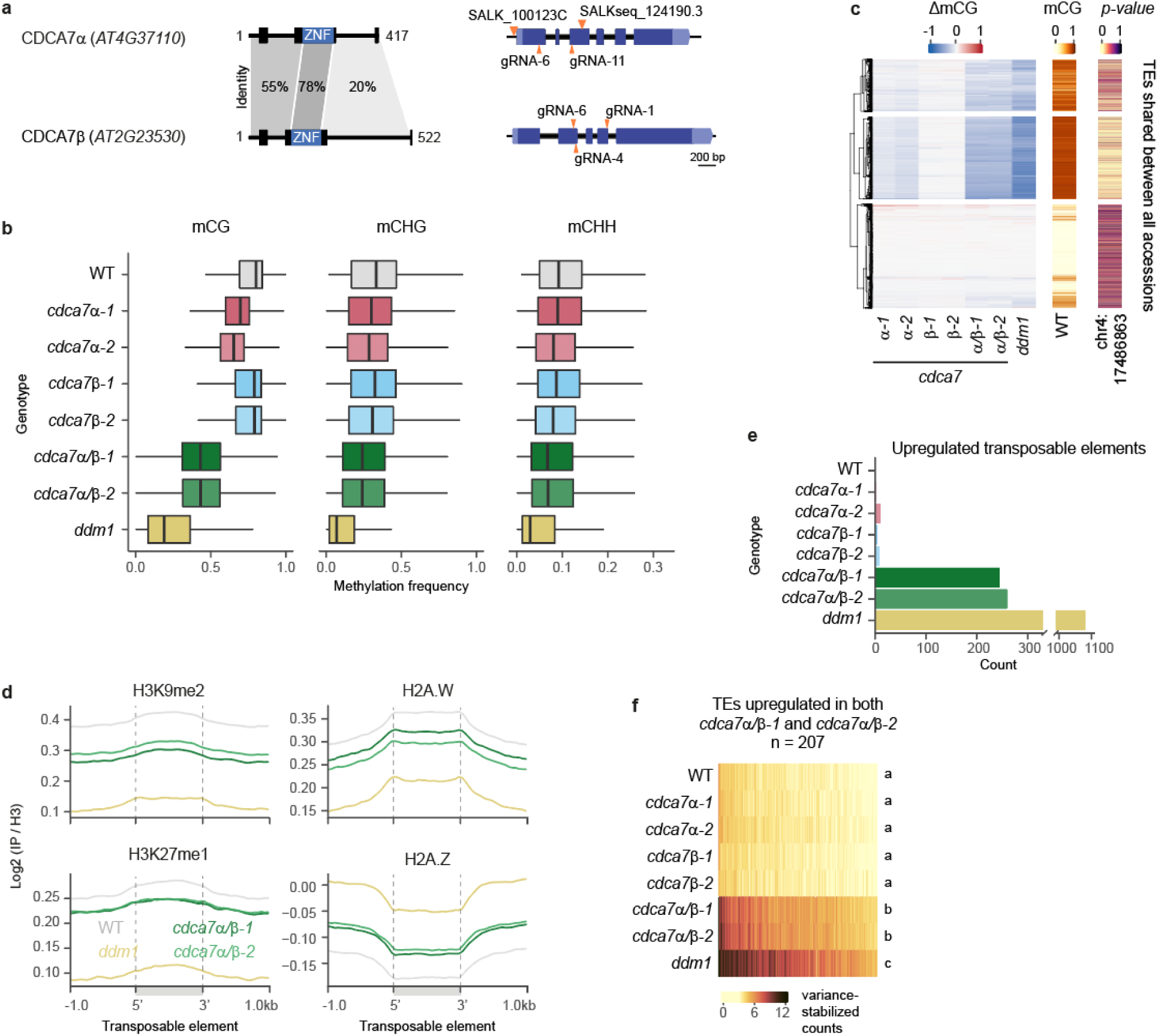
CDCA7α and CDCA7β redundantly maintain mCG, heterochromatin marks and transcriptional silencing. **a,** Protein (left) and gene (right) models for CDCA7α and CDCA7β, with sequence identity scores between domains. Secondary structures (boxes) and low confidence regions (black lines, pLDDT < 70) are shown. Orange triangles mark T-DNA insertions and CRISPR-Cas9 gRNAs. **b,** 5mC in indicated contexts at TEs. **c**, Heatmap comparing ΔmCG (mutant - WT) for *cdca7* null mutations with the effect of *CDCA7α* natural alleles on mCG, using the association of the SNP at chr4:17486863 with TE mCG (*p-value*). K-mean clusters were defined based on ΔmCG. The WT value is shown as a reference. **d,** Metaplots of mean levels of histone marks and variants at heterochromatic TEs. **e,** TEs upregulated in the indicated mutants relative to WT. **f,** Transcript levels of TEs upregulated in both *cdca7α/β* mutants, shown with variance-stabilized counts. Lowercase letters denote statistical groups determined by Tukey’s HSD test (*P* < 0.05).

To investigate the functions of these CDCA7 paralogs, we generated single and double mutants for *cdca7α* and *cdca7β* using T-DNA insertions and CRISPR-Cas9, obtaining two independent lines per mutant to control for potential genetic aberrations (**Fig. 2a, S5a**). We compared them with *ddm1* mutants of the same generation (see methods), because the defects caused by *ddm1* loss of function gradually worsen over consecutive homozygous generations (Kakutani *et al*., 1996; Ito *et al*., 2015).

TEs had reduced mCG levels in the two *cdca7α* mutant alleles, with a stronger effect in *cdca7α-2* (**Fig. 2b**). The T-DNA in *cdca7α-2* disrupts the coding sequence, whereas the T-DNA in *cdca7α-1* is located in the promoter (**Fig. S5a**). Single mutations in *CDCA7β* did not affect mCG levels, but *cdca7α/β* double mutants had strongly decreased mCG compared to *cdca7α* (**Fig. 2b**), demonstrating that *CDCA7α* and *CDCA7β* redundantly maintain mCG levels. The *cdca7α* and *cdca7β* mutations had only minimal impact on mCHG and mCHH levels, with a synergistic effect in *cdca7α/β* (**Fig. 2b**). Compared with *cdca7α/β*, loss of DDM1 had a stronger impact in all contexts (**Fig. 2b**). Mirroring our GWAS results, the impact of *cdca7α/β* was strongest at CMT2-dependent TEs (**Fig. S5c**). Importantly, TEs with the strongest mCG decrease in *cdca7α* and *cdca7α/β* also had the most significant association of the *CDCA7α* SNP with mCG variation at CMT2-dependent TEs (**Fig. 2c**), indicating that null and natural alleles of *CDCAα* impact the same targets.

Because DDM1 directly controls the genomic distribution of histone H2A.W and H2A.Z, and affects the H3K9me2 and H3K27me1 histone post-translational modifications (Soppe *et al*., 2002; Ikeda *et al*., 2017; Osakabe *et al*., 2021; Jamge *et al*., 2023; Lee *et al*., 2023), we profiled these chromatin marks in *ddm1* and two *cdca7α/β* mutants using ChIP-seq, focusing on heterochromatic TEs (see methods). Our results showed that *cdca7α/β* mutants exhibited decreased levels of H3K9me2, H3K27me1 and H2A.W, while H2A.Z levels increased, resembling *ddm1* but with milder effect (**Fig. 2d**).

Furthermore, we quantified the impact of *cdca7* mutations on TE silencing using 3’-Tag RNA-seq. Single mutations in *cdca7α* or *cdca7β* had negligible effects, but the double mutants showed over 200 upregulated TEs (**Fig. 2e**). The extent of TE upregulation in *cdca7α/β* was less than in *ddm1* mutants (**Fig. 2f**). To further validate these observations, *cdca7α/β-1* was complemented with *CDCA7α* or *CDCA7β*, fused with an N or C-terminal tag. Constructs with a C-terminal tag partially complemented silencing defects (**Fig. S5d**), confirming a direct role of *CDCA7α*/*β* in TE silencing. Interestingly, the N-terminal tag inhibited *CDCA7α*/*β*-mediated silencing, suggesting it interfered with a critical CDCA7 function.

We found no developmental defects in *cdca7α/β* mutants and, in contrast to *ddm1* (Kakutani *et al*., 1996; Ito *et al*., 2015), inbreeding homozygous mutants for up to eight generations had no evident effect on the plant phenotype (**Fig. S5e**). In summary, *CDCA7α* and *CDCA7β* redundantly maintain heterochromatin marks and TE silencing. *CDCA7α* and *CDCA7β* appear to have identical functions, but *CDCA7α* plays a more prominent role than *CDCA7β* (**Fig. 2b**), possibly due to its higher expression (**Fig. S4**).

### CDCA7α and CDCA7β require DDM1 to maintain CG methylation

At the molecular level, *cdca7α/β* null mutants resembled *ddm1* (**Fig. 2)**, suggesting CDCA7α/β and DDM1 operate in a common pathway. To test this genetically, we combined *ddm1* and *cdca7α/β* mutations and measured their impact on 5mC and TE silencing. The triple *cdca7α/β ddm1* mutants showed 5mC levels comparable to *ddm1* mutants of the same generation (**Fig. 3a, b**). Additionally, introducing any *cdca7* mutation alongside *ddm1* did not alter TE transcript levels (**Fig. 3c, S6a**). Therefore, we conclude that CDCA7α/β fully operate through DDM1 to regulate 5mC and TE silencing.

**Figure 3.**
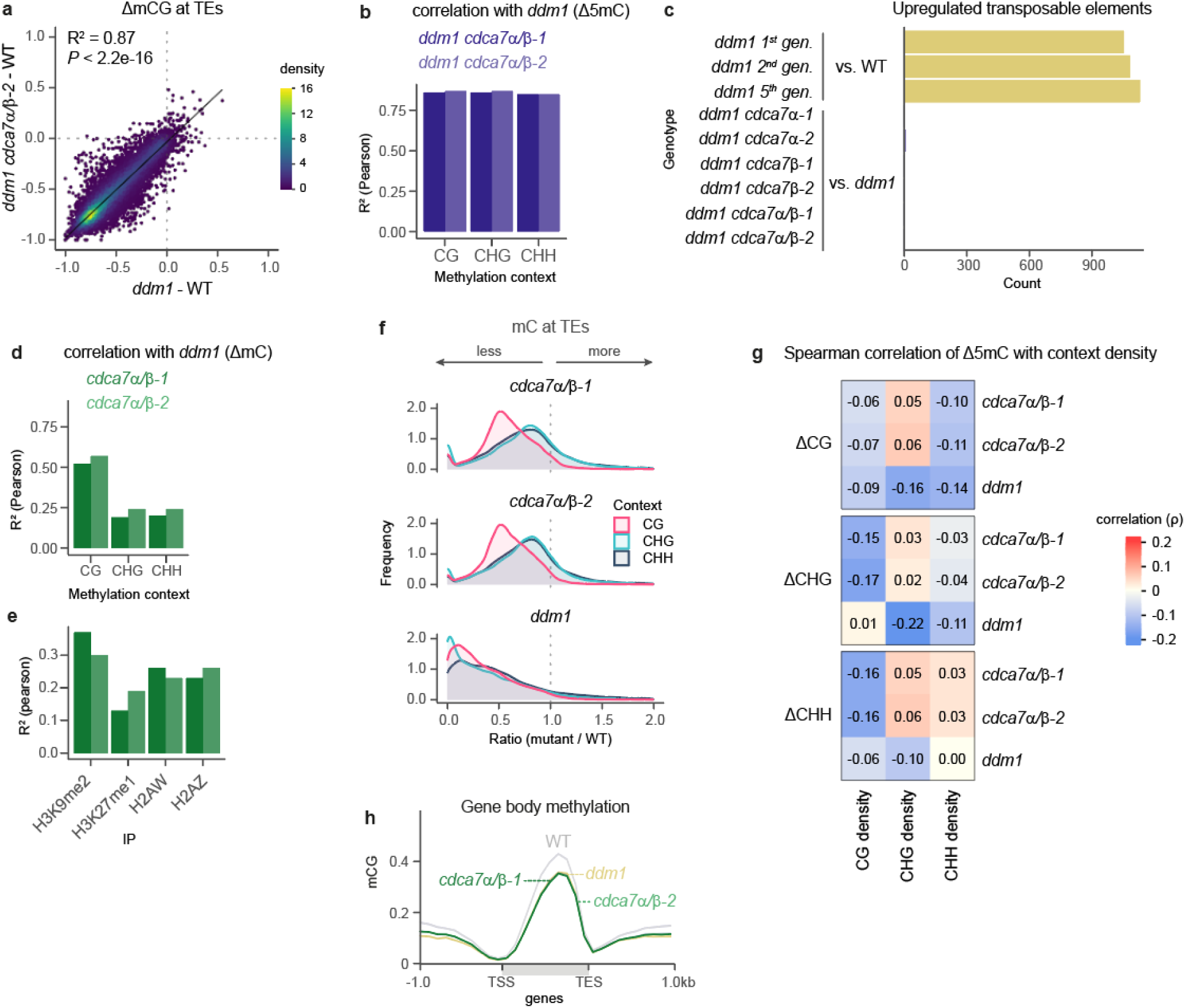
CDCA7α and CDCA7β function through DDM1 to maintain mCG. **a**, Linear correlation of mCG loss at TEs in *ddm1 cdca7α/β-2* compared to *ddm1*. Pearson correlation coefficients (R^2^) and a permutation test-derived *P*-value are indicated. **b,** Pearson correlation coefficients (R^2^) of TE 5mC loss in *ddm1* and the indicated mutants. All correlations had *P* < 1e-10. **c**, Number of transposable elements upregulated in the indicated mutants relative to WT or *ddm1*. Mutant combinations of *ddm1* and *cdca7* were compared to *ddm1* mutants of the corresponding generation. **d**, **e**, Pearson correlation coefficients (R^2^) showing the relationship between ddm1 and the indicated mutants in terms of 5mC loss (**d**) and chromatin mark changes (**e**). All correlations had *P* < 1e-10. **f**, Distribution of methylation ratios (mutant / WT) across genotypes and 5mC contexts. The x-axis is capped at 2, excluding at most 3.2% of the data. **g**, Spearman correlations coefficients (ρ) of methylation context density (number of CN sites per bin) with 5mC loss in mutants, calculated in 100 bp bins. **h**, Metaplot showing mean gene body methylation in CG context. Only gene-body methylated genes are included (n=9938).

In all assays, loss of CDCA7α/β had a milder effect than the loss of DDM1 (**Fig. 2**), suggesting that CDCA7α/β might specifically target a subset of DDM1-dependent TEs. To test this hypothesis, we compared TEs upregulated in both mutants. Although TEs upregulated in *cdca7α/β* were a subgroup of those upregulated in *ddm1* (**Fig. S6b**), 51.8% of TEs upregulated exclusively in *ddm1* had increased transcript levels in *cdca7α/β*, but did not meet significance thresholds (**Fig. S6c**). Additionally, all TEs silenced by DDM1 showed reduced 5mC in *cdca7α/β*, irrespective of their expression levels (**Fig. S6d**). This demonstrates that *CDCA7α/β* is required for the maintenance of 5mC at all TEs silenced by *DDM1*, not just a specific subset. We extended the analysis to all TEs with mCG in the WT and found that 91.4% of these lost mCG in both *cdca7α/β-1* and *cdca7α/β-2* (**Fig. S6d**), demonstrating that CDCA7α/β target nearly all TEs.

DDM1 has two distinct functions in heterochromatin maintenance: first, depositing H2A.W while preventing deposition of H2A.Z (Osakabe *et al*., 2021; Jamge *et al*., 2023; Zhou *et al*., 2023) and second, facilitating access of DNA methyltransferase to nucleosomal DNA through nucleosome remodeling (Zemach *et al*., 2013; Lyons and Zilberman, 2017). Heterochromatic histone modifications also depend on DDM1 (Soppe *et al*., 2002; Ikeda *et al*., 2017; Osakabe *et al*., 2021; Lee *et al*., 2023) through unknown mechanisms. CDCA7α/β might therefore differentially affect specific functions of DDM1. To explore this, we compared the effects of *ddm1* and *cdca7α/β* mutations on 5mC, histone variants and modifications. The effects of the mutations correlated strongly for mCG (**Fig. 3d**, **S6f**, R^2^=0.52-0.57) but weakly for mCH (R^2^=0.19-0.24). Relative to these, correlations were intermediate for H3K9me2 (**Fig. 3e**, R^2^=0.30-0.37), likely due to H3K9me2 being reliant on both mCH (Du *et al*., 2015) and mCG (Tariq *et al*., 2003; Mathieu *et al*., 2007; Deleris *et al*., 2012; Choi, Lyons and Zilberman, 2021). For H3K27me1 and histone variants H2A.W and H2A.Z, correlations were low (**Fig. 3e**), although overall trends resembled those seen in *ddm1* (**Fig. S6g**). Therefore, *cdca7α/β* mutants primarily resembled *ddm1* in their effect on mCG and not on deposition of histone variants or histone post-translational modification.

Strikingly, *cdca7α/β* mutants were disproportionately affected in CG context, whereas all 5mC contexts were equally affected in *ddm1* (**Fig. 3f**), highlighting the specific role of CDCA7α/β in regulating mCG. Likewise, GWAS pointed towards a distinct role of CDCA7α/β in mCG regulation (**Fig. S1**). Finally, and in contrast with the mild effect on other marks, loss of CDCA7α/β accounted for 60% of the mCG changes observed in *ddm1* mutants (**Fig. S6h**). Taken together, these data demonstrate that CDCA7α/β are primarily involved in mCG maintenance.

In *met1* mutants, mCG loss leads to depletion of mCHG, mCHH, and H3K9me2, with a concurrent increase in H2A.Z (Soppe *et al*., 2002; Tariq *et al*., 2003; Zilberman *et al*., 2008; Stroud *et al*., 2013; Choi, Lyons and Zilberman, 2021). This suggests that the moderate changes in levels of mCH, H2A variants and H3 modifications in *cdca7α/β* mutants are secondary effects of reduced mCG. In support of this interpretation, we found that the changes of mCHG and mCHH in *cdca7α/β* mutants was anticorrelated with the density of CG sites (**Fig. 3g**). This differed from *ddm1*, where the loss of mCH was best correlated with CHG density.

We next examined gene bodies, where mCG is decoupled from mCH (Muyle *et al*., 2022). The absence of CDCA7α/β mirrored the impact of *ddm1* mutations (**Fig. 3h**), indicating that CDCA7α/β is fully required for DDM1-dependent gene body methylation. This is in accordance with a model in which CDCA7α/β specifically regulates mCG. We conclude that CDCA7α/β primarily regulate DDM1’s function in maintaining mCG without directly affecting its other roles.

### CDCA7 interacts with DDM1 through conserved regions

Next, we explored the function of the three CDCA7α/β domains using a genetic complementation strategy. The N-terminal domain contains a highly conserved motif (**Fig. 4a**), the 4CXXC zinc finger domain binds hemimethylated CG dinucleotides (Hardikar *et al*., 2024; Shinkai *et al*., 2024; Wassing *et al*., 2024) and the C-terminal part of the protein has lower conservation.

**Fig 4.**
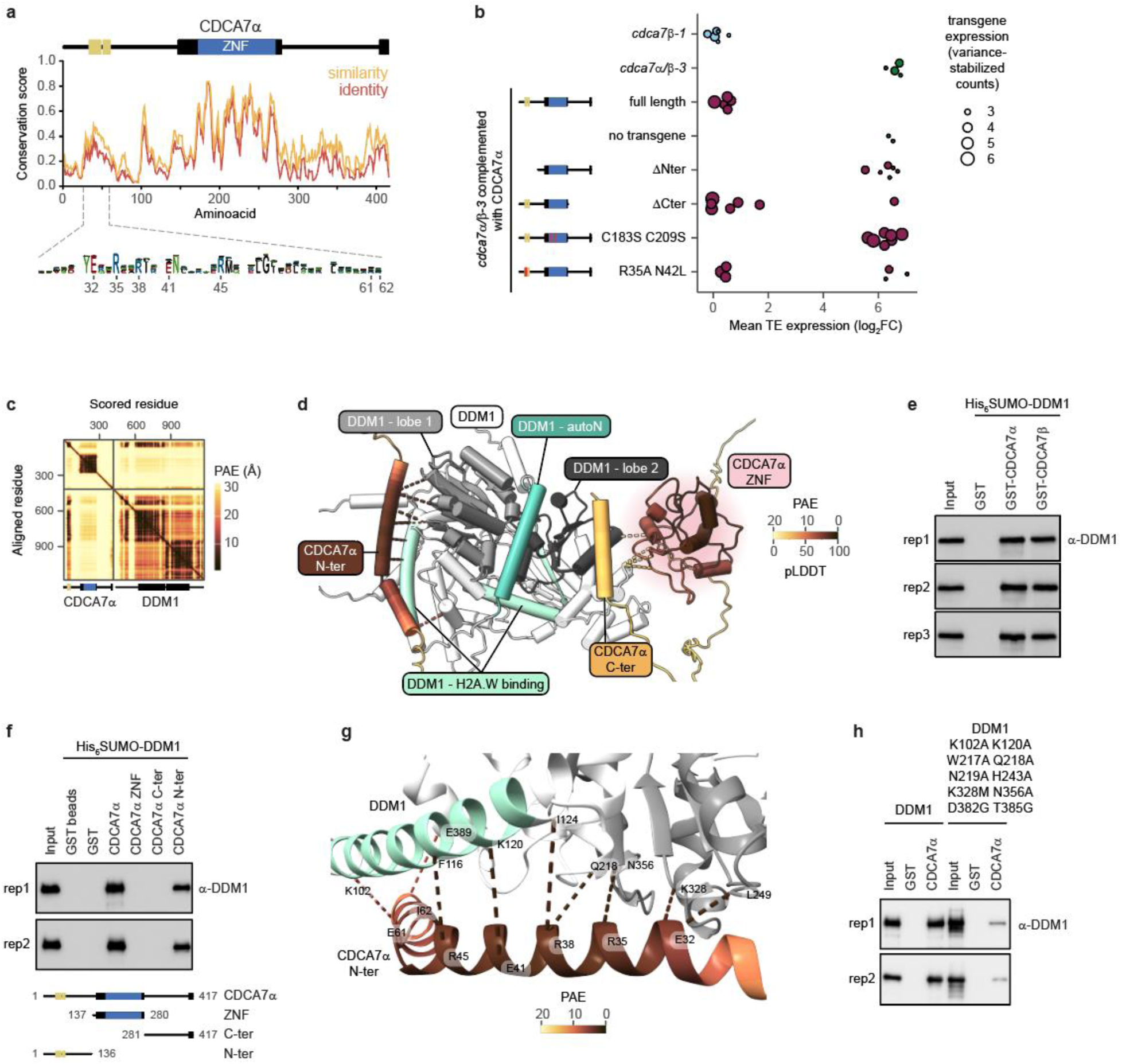
Characterization of CDCA7α/β protein domains and DDM1 interaction. **a**, Conservation of CDCA7α among 315 orthologs in Viridiplantae, with domains represented to scale, including the DDM1-interacting alpha-helices (yellow). A consensus sequence (bottom, n = 37) highlights residues predicted to contact DDM1. **b**, TE upregulation in *cdca7α/β-3* complemented with various CDCA7α constructs. Each data point is a T1 plant, showing the mean log_2_ fold change (relative to WT) of TEs upregulated in *cdca7α/β-3*, scaled by transgene expression. **c**, Predicted Aligned Error (PAE) of the modeled interaction between CDCA7α and DDM1, with protein domains indicated. **d**, Predicted CDCA7α-DDM1 structure, showing predicted contacts ≤ 3 Å colored by PAE. CDCA7α is colored by local confidence, using predicted local distance difference test (pLDDT). **e**, In vitro co-immunoprecipitation (Co-IP) of recombinant CDCA7α and CDCA7β with DDM1, detected by Western Blot. **f**, Co-IP of recombinant DDM1 with truncated CDCA7α constructs. **g**, Predicted interaction of CDCA7α N-terminal alpha helices with DDM1. Contacts ≤ 3 Å are shown, colored by PAE. **h**, Co-IP of recombinant wild-type and mutant DDM1 proteins with CDCA7α.

Once mCG is lost in *met1* or *ddm1* mutants, it can largely not be restored by complementation (Finnegan, Peacock and Dennis, 1996; Kakutani *et al*., 1999; Lippman *et al*., 2003; Saze, Scheid and Paszkowski, 2003), likely because of the semi-conservative nature of mCG maintenance. Accordingly, complementation of *cdca7α/β-1* was partial and locus-specific (**Fig. S5d**). To bypass this limitation, we introduced a *CDCA7α* transgene before the complete loss of endogenous *CDCA7α/β* (**Fig. S7a, b**). We quantified TE transcripts in primary transformants using transcriptomes and took into account transgene expression, since it is highly variable between primary transformants (Matzke and Matzke, 1998). Untransformed *cdca7β-1* and *cdca7α/β-3* mutants showed no or few reads mapping to the transgenic tag, reflecting background noise (**Fig. 4b**). As expected, a full length CDCA7α transgene restored TE silencing, while sister plants lacking the transgene had elevated TE transcript levels. CDCA7α(ΔC-ter) complemented silencing defects in individuals with high transgene expression, indicating that the C-terminal domain is dispensable for TE silencing. Constructs with an N-terminal deletion were not expressed, preventing conclusions on the role of the N-terminus in silencing (**Fig. 4b**). To test the function of the zinc finger domain, we mutated the conserved cysteines C183 and C209, which coordinate a zinc ion (**Fig. S8a**). The lack of complementation with this construct (**Fig. 4b**) indicates that the CDCA7α zinc finger is essential for TE silencing.

To further explore the function of CDCA7α/β domains, we used AlphaFold-Multimer (Evans *et al*., 2022) to model potential interactions with DDM1. High confidence interactions were identified between CDCA7α/β N-termini and DDM1, specifically its lobe 1 and H2A.W binding domain (**Fig. 4c-d, S8b**). In vitro immunoprecipitation assays with purified recombinant proteins showed that both CDCA7α and CDCA7β bind to DDM1 (**Fig. 4e**), and these interactions were maintained under more restrictive salt concentration (**Fig. S8c**). The N-terminal region of CDCA7α alone was sufficient to interact with DDM1 (**Fig. 4f**), whereas neither the Zn finger nor the C-terminal part showed an interaction. Remarkably, the N-terminal residues predicted to interact with DDM1 (**Fig. 4g**) were highly conserved across the green lineage (**Fig. 4a**). CDCA7α R35 and N42 were also conserved in model vertebrates and predicted to interact with HELLS (DDM1 ortholog) (**Fig. S8d**). However, R35A and N42L mutations did not disrupt binding to DDM1 (**Fig. S8e**) or CDCA7α silencing function (**Fig. 4b**), indicating that other CDCA7α residues can mediate DDM1 binding in *A. thaliana*. DDM1 residues predicted to interact with CDCA7α were also conserved (**Fig. S8f**), and their mutation reduced the interaction with CDCA7α (**Fig. 4h**). Since DDM1 activity is restricted by a N-terminal autoinhibitory coiled-coil domain (autoN) (Lee *et al*., 2023; Nartey, Goodarzi and Williams, 2023), we considered that CDCA7α binding could stimulate DDM1 by displacing the autoN. However, the position of the autoN was predicted with poor confidence relative to DDM1 lobes (**Fig. S8g**) and was not improved by co-modeling CDCA7a/b or a nucleosome (**Fig. 4c, S8h, i**), providing no evidence that CDCA7α stimulates DDM1 activity through autoN displacement. Further modeling suggested that CDCA7α/β paralogs could not bind to each other or themselves, and that DDM1 can bind only one CDCA7α/β paralog at a time (**Fig. S8j**), consistent with CDCA7α/β acting redundantly (**Fig. 2**). We conclude that the conserved N-terminal domain of CDCA7α/β directly interacts with DDM1 *in vitro*.

In summary, the C-terminal domain of CDCA7α does not contribute to its silencing function, while the zinc finger is essential. In both CDCA7α and vertebrate CDCA7, the zinc finger binds hemi-methylated CG (Hardikar *et al*., 2024; Shinkai *et al*., 2024; Wassing *et al*., 2024), implying that CDCA7α recruits DDM1 to hemi-methylated CG sites, facilitating mCG deposition after DNA synthesis. This recruitment is mediated by a conserved N-terminal domain of CDCA7, which also interacts with HELLS in vertebrates (Wassing *et al*., 2024). Combined with our previous results, our biochemical data indicate that DDM1 interacts directly with either CDCA7α or CDCA7β, and this interaction is crucial for maintaining mCG and repressing TEs.

### Genetic variation in the *CDCA7*α promoter shapes large mCG variation

Next, we investigated the molecular causes of the *CDCA7α* natural alleles. Although protein sequences showed a cysteine to serine mutation at position 182, this cysteine is not conserved in sister species (**Fig. S9**), nor in eukaryotes (Funabiki *et al*., 2023), and it does not coordinate a zinc atom (**Fig. S8a**). There were no amino acid changes in conserved sites within the zinc finger domain or the DDM1-interacting domain, suggesting that a causal effect on protein activity was unlikely (**Fig. S9**). Therefore, to explore the impact of natural genetic variation on *CDCA7α* regulation, we performed GWAS for *CDCA7α* expression using published transcriptome data (*n* = 427 lines; (Kawakatsu *et al*., 2016; Kornienko *et al*., 2023)).

Remarkably, a strong *cis* effect was detected for *CDCA7α* expression, with the most significant SNP identified as chr4:17486863 (*CDCA7α-alt_a_* haplotype) (*p*-value=7.8E-43, MAF=0.33), the same allele identified in our GWAS for mCG levels (**Fig. 1a-c, 5a**). Additionally, chr4:17497441_non-ref_ (*CDCA7α-alt_b_* haplotype) was also present in this large peak (*p*-value=2.08E-6, MAF=0.06). Both haplotypes were associated with higher *CDCA7α* expression, two to three times higher than the reference allele Col-0, as well as elevated mCG levels (**Fig. 5b**).

**Figure 5.**
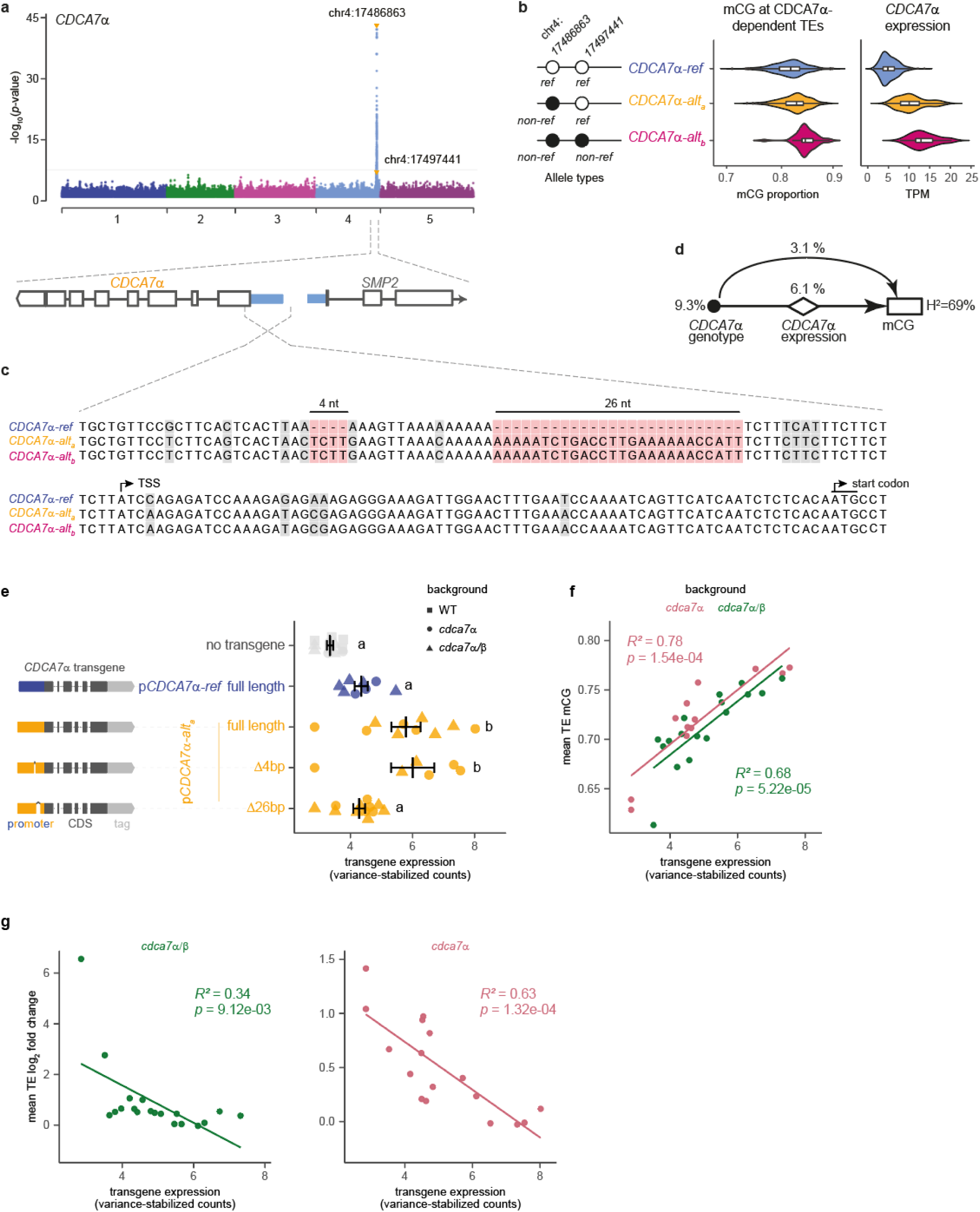
Cis-polymorphism regulates CDCA7α promoter activity and shapes mCG variation. **a**, GWAS for *CDCA7*α expression. SNPs identified in GWAS for mCG are highlighted with orange dots. **b**, The allelic effects of *CDCA7*α on mCG and the expression levels. Violin plots show mean mCG levels of CDCA7α-targeted TEs (*n*=774 lines) and *CDCA7*α expression for each genotype (*n*=461 lines). **c**, Sequence alignment upstream of *CDCA7a* in three haplotypes. **d**, A mediation model shows the genetic effect of *CDCA7*α alleles on mCG through *CDCA7a* expression. Black plot, diamond, and rectangle indicate genotype, gene expression, and mCG phenotype, respectively. Arrows show the regulation. **e**, Effect of CDCA7α promoters on transgene expression, measured by mRNA-seq in complemented *cdca7α/β-3* and *cdca7α-2* mutants. Transgenic constructs are shown to the left (to scale, except deletions). Each data point is an individual primary transformant with mean values ± standard error shown in black and significance between groups indicated by lowercase letters (one-way ANOVA with Tukey’s post-hoc tests, *p* < 0.05). **f**,**g**, Correlation of transgene expression with mean TE CG methylation (**f**) or mean TE log_2_(fold change), relative to WT (**g**). Pearson correlation coefficients and *p*-values (permutation test) are shown.

To test for a causal link between DNA methylation and *CDCA7α* expression, we performed a mediation analysis (Sasaki, Frommlet and Nordborg, 2018). This showed that 65.6% of the *CDCA7α* genetic effect was mediated by *CDCA7α* expression, suggesting that overexpression of *CDCA7α* in *CDCA7α-alt_a_* and *CDCA7α-alt_b_* haplotypes contributes to the divergence of mCG levels (**Fig. 5d**). Alignments of the *CDCA7α* promoter region using fully assembled genomes revealed that the region is unusually polymorphic. Two indels distinguished *CDCA7α-alt_a_* and *CDCA7α-alt_b_* from the reference-type haplotype (*CDCA7α-ref*) (**Fig. 5c**). This suggests that different haplotypes in the *CDCA7α* promoter region change *CDCA7α* expression, resulting in diverse mCG levels in wild populations of *A. thaliana*.

To test the regulatory effect of the *CDCA7α* haplotypes on *CDCA7* expression, we cloned the *CDCA7α-alt_a_* promoter and fused it to the *CDCA7α-ref* coding sequence to complement *cdca7α* and *cdca7α/β-3* mutants (**Fig. S7a**). We also generated constructs where the indels present in *CDCA7α-alt_a_* and *CDCA7α-alt_b_* were deleted. In primary transformants, transgene expression was higher with the *CDCA7α-alt_a_* promoter, and this effect depended on the presence of a 26 bp indel but not a 4 bp indel (**Fig. 5e**). Importantly, *CDCA7α* expression positively correlated with mCG levels at TEs (**Fig. 5f**) and inversely correlated with TE expression (**Fig. 5g**), regardless of the mutant background. These findings demonstrate that the 26 bp indel in the *CDCA7α* promoter regulates its expression, which in turn controls DNA methylation levels in natural *A. thaliana* populations. The fact that higher expression of *CDCA7α* led to increased mCG levels further implies that in nature, *CDCA7α* is a limiting factor in promoting mCG and TE repression.

The construct lacking the 26 bp indel, despite producing *CDCA7α* transcript levels similar to those of the reference promoter (**Fig. 5e**), was associated with reduced mCG (**Fig. S10a**), suggesting that steady-state *CDCA7α* transcript levels are not sufficient to predict mCG levels. There are six reported *CDCA7α* transcript isoforms (Zhang *et al*., 2022). Under our experimental conditions, we detected three: *CDCA7α.2*, *CDCA7α.3*, and *CDCA7α.5* (**Fig. S10b**). *CDCA7α.2*, the most abundant isoform (**Fig. S10b**), encodes the canonical CDCA7α protein, while *CDCA7α.3* has no annotated protein product and *CDCA7α.5* produces a protein lacking the DDM1-interacting domain. We found that both *CDCA7α.2* and *CDCA7α.3* were overexpressed in lines with the *CDCA7α-alt_a_* promoter, and this overexpression required the 26 bp indel (**Fig. S10c**), consistent with our quantification of all isoforms (**Fig. 5e**). However, deleting the 26 bp indel had no strong effect on the *CDCA7α.5* isoform relative to the intact *CDCA7α-alt_a_* promoter (**Fig. S10c**), indicating that this indel regulates isoform usage. *CDCA7α.5*, which lacks the DDM1-interacting domain but retains an intact zinc finger, could compete with the canonical CDCA7α for binding to target loci, thereby antagonizing DDM1 activity. We hypothesize that an increased abundance of *CDCA7α.5*, along with reduced levels of *CDCA7α.2*, would contribute to the observed reduction in mCG levels in transgenic lines with *CDCA7α-alt_a_* promoters lacking the 26 bp indel (**Fig. S10a**).

In conclusion, the three *CDCA7α* haplotypes influence DNA methylation and TE repression through different mechanisms. The *CDCA7α-alt_a_* and *CDCA7α-alt_b_* haplotypes, with their unique promoter sequence, drive higher CDCA7α expression and corresponding mCG levels compared to the reference haplotype. Moreover, the 26 bp indel plays a critical role not only in regulating *CDCA7α* expression but also in controlling the balance of transcript isoforms, which has downstream effects on mCG levels. While no genetic difference between *CDCA7α-alt_a_* and *CDCA7α-alt_b_* haplotypes was observed in the tested promoter region, additional structural variants that could explain their expression variation likely exist.

### The *CDCA7α*-ref allele is derived and predominantly occurs in Europe

We investigated the history of the *CDCA7α* promoter region to gain insight into the evolution of the epigenome in *A. thaliana*. To understand which form of the *CDCA7α* promoter is ancestral, we compared the *CDCA7α* haplotypes with closely related species, including *A. lyrata*, *A. halleri*, and *Capsella rubella*. These species clearly exhibited higher similarity to the *CDCA7α-alt_a_* and *CDCA7α-alt_b_* haplotypes than to *CDCA7α-ref* (**Fig. 6a**). The predicted haplotype network supported the conclusion that the *CDCA7α-alt_a_* haplotype, causing higher mCG levels, is ancestral, while *CDCA7α-ref* is derived, and the promoter indels reflect deletions (**Fig. 6b**). Compared with *CDCA7α-alt_a_*, *CDCA7α-ref* has a 12.3% shorter promoter region (**Fig. 6c**). *CDCA7α-alt_b_*, which drives the highest mCG levels, accumulated specific SNPs not shared with relative species, and the haplotype network showed that *CDCA7α-alt_b_* was derived from *CDCA7α-alt_a_* (**Fig. 6b**). Consequently, the regulation of *CDCA7α* expression has undergone multiple changes, resulting in variation in mCG levels at TEs in heterochromatic regions.

**Figure 6.**
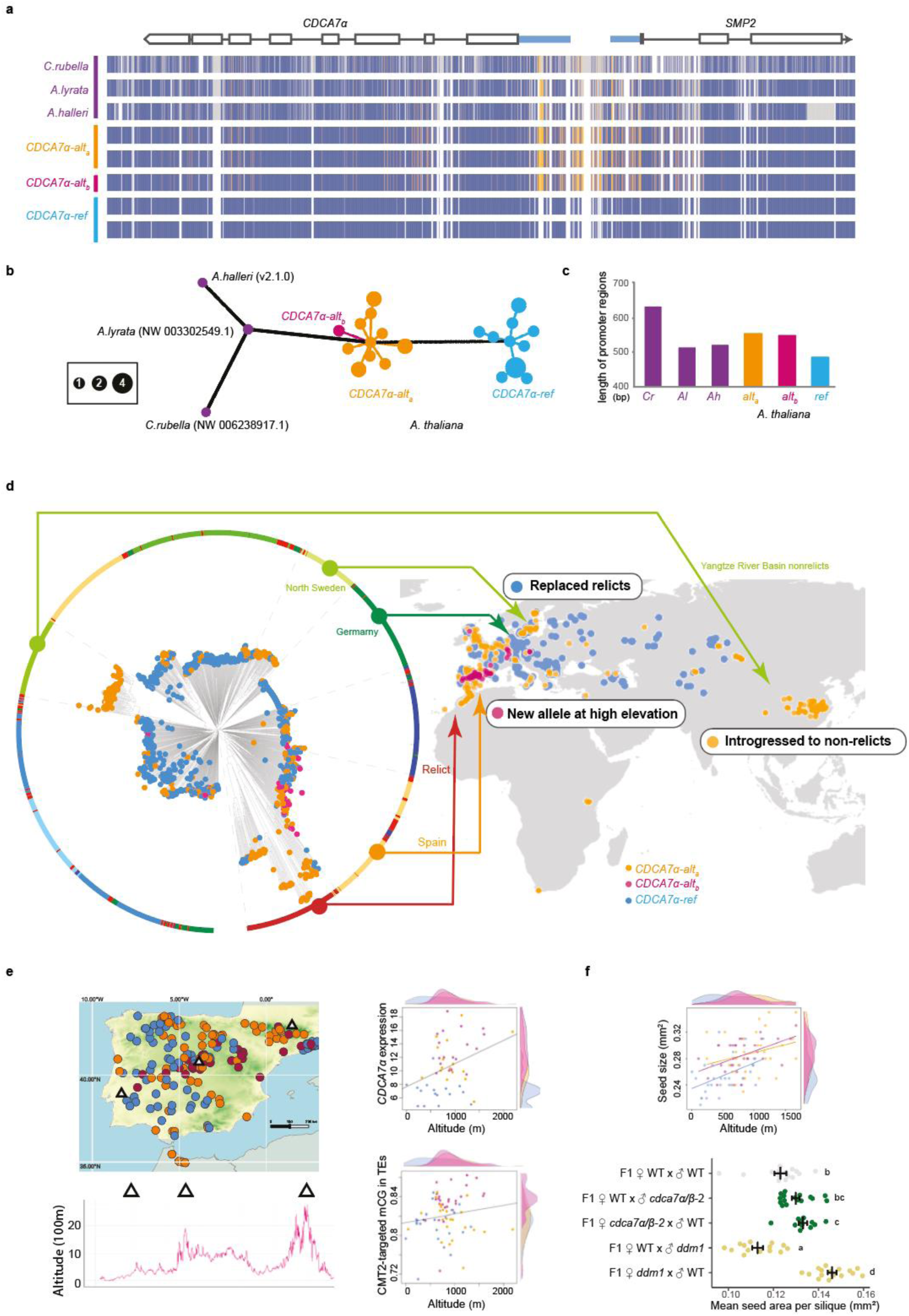
The evolutionary history of *CDCA7*α haplotypes. **a,** Alignment of *CDCA7*α regions and the ancestral state. Blue, orange, and gray regions represent the reference, the alternative, and non-*A. thaliana* alleles, respectively. **b**, Haplotype network of a 4 kbp region covering the *CDCA7*α and *SMP2* coding region. **c**, Variation in *CDCA7*α promoter length. **d,** Population structures of 1323 natural lines with *CDCA7*α alleles and the geographic distribution. The phylogenetic tree is a neighbor-joining tree (Hsu et al., 2019) with populations (outer circle) and the genotype of *CDCA7*α haplotypes (at the tips). A global map shows an overview of allele distributions. **e**, Altitudinal cline of alleles and mCG variation. The local map of the Iberian Peninsula indicates the altitudes of summits with triangles, and scatterplots show associations between altitude and *CDCA7*α expression (top) and mCG in CMT2-targeted TEs (bottom), with the allele distribution as density plots. **f**, The effect of *CDCA7*α on seed size. A scatter plot shows the association between altitude and seed size, with the allele distribution shown as density plots (reanalyzed data from Vidigal et al. 2016). Dot plots (bottom) show parent-of-origin effects of *cdca7α/β-2* and *ddm1* on seed size, in F1 crosses to WT. Lowercase letters denote statistical groups determined by Tukey’s HSD test (*P* < 0.05).

The *CDCA7α-alt_a_* haplotype was predominantly found in three regions: (1) Northern Sweden, (2) the Iberian peninsula, and (3) Eastern Asia, specifically the Yangtze River Basin population, which is the most southern-eastern edge of native *A. thaliana* habitats (**Fig. 6d**). Consistent with *CDCA7α-alt_a_* being ancestral, its distribution overlaps with the ancestral-type populations named ‘relict’ observed in Northern Sweden and the Iberian peninsula (1001 Genomes Consortium, 2016; Lee *et al*., 2017). Relicts have genomes that diverged from those of the current worldwide population, referred to as ‘non-relicts,’ and are considered to have been dominant in post-glacial Eurasia (1001 Genomes Consortium, 2016; Lee *et al*., 2017). Non-relicts comprising over 95% of current populations are likely modern populations that replaced the relicts as they spread to central and western Europe through human activity. Most of the lines carrying the *CDCA7α-ref* were non-relicts (**Fig. 6d**). Although the Yangtze River Basin population is non-relict, expected to have diverged about 60 thousand years ago (Hsu, Lo and Lee, 2019), *CDCA7α-alt_a_* was fixed in this relatively new population (**Fig. 6d**). This is in agreement with studies suggesting that genomic introgression occurred from local relicts to the Yangtze River Basin population (Hsu, Lo and Lee, 2019), and the *CDCA7α* region appears to have been introgressed as shown in our phylogenetic tree (**Fig. 6d**). Additionally, a prior selection scan using the composite likelihood ratio (CLR) in Zou et al (2017) indicated a signature of positive selection over a 15 kbp region that includes *CDCA7α*, suggesting that this allele may be involved in local adaptation.

Unlike *CDCA7α-alt_a_*, *CDCA7α-alt_b_* was predominantly found in the Spanish population, particularly in the central mountain region of the Iberian Peninsula (**Fig. 6d, e**). This mountainous area stretches from east to west, exhibiting diverse climates due to its wide range of altitudes, up to 3000 m above sea level (Vidigal *et al*., 2016). While *CDCA7α-alt_b_* appears to have been derived from *CDCA7α-alt_a_*, non-relict Spanish populations carry *CDCA7α-alt_b_* (**Fig. 6d**), suggesting that *CDCA7α-alt_b_* likely emerged after *CDCA7α-alt_a_* was introgressed into non-relicts. Across the Spanish population in the Iberian Peninsula, both *CDCA7α-alt_a_* and *CDCA7α-alt_b_* were distributed at higher altitudes than *CDCA7α-ref*, resulting in an altitudinal cline of mCG levels on TEs (**Fig. 6e**).

If *CDCA7α* is directly involved in expansion to new environments, *CDCA7α*-mediated epigenetic changes must impact life-history traits that provide adaptive features. For example, flowering time and seed size are influenced by DNA methylation (Adams *et al*., 2000; Kinoshita, 2004; Xiao *et al*., 2006) and contribute to reproductive success in challenging environments (Alonso-Blanco *et al*., 1999; Gaudinier and Blackman, 2020). While *cdca7α/β* mutants did not affect flowering time under laboratory conditions (**Fig. S12**), a maternal effect was observed in seed size (**Fig. 6f**). This differed from *ddm1*, which showed the expected parent-of-origin effect (Adams *et al*., 2000; Xiao *et al*., 2006; FitzGerald *et al*., 2008). Consistent with the mutant phenotype, seed size variation in the Spanish mountain population showed a clear altitudinal cline and was positively associated with the distribution of *CDCA7α* alleles along altitude (**Fig. 6f**). While seed size is a complex trait controlled by multiple loci (Alonso-Blanco *et al*., 1999; Gnan, Priest and Kover, 2014), our results suggest mCG changes through *CDCA7α* alleles potentially contribute, and may play a role in colonizing new environments.

## Discussion

The establishment and maintenance of the epigenome, particularly mCG, a DNA modification that is transgenerationally inherited, has been much debated (Richards, 2006; Charlesworth, Barton and Charlesworth, 2017; Baduel and Sasaki, 2023). Our study provides novel molecular mechanisms that explain the epigenome variation observed in nature. We identified *CDCA7α* as a new *trans* regulator of mCG in plants and showed that its expression variation has strongly shaped the epigenome of natural populations in *A. thaliana*. The ancestral *CDCA7α* promoter allele, present in relict populations, drives higher *CDCA7α* expression compared to the derived alleles found in non-relict populations across western and central Europe, including the reference strain Col-0. A 26 bp deletion in the derived promoter reduces *CDCA7α* expression and increases the production of a truncated *CDCA7α* isoform, with the potential to inhibit DDM1 activity. Together, these changes result in lower mCG levels and reduced TE repression in current worldwide populations compared to the ancestral form.

In addition to natural variation in *CDCA7α,* the retention of two *CDCA7* copies following whole genome duplication (**Fig. S3**) may have influenced mCG dynamics. In jawed vertebrates, *CDCA7* is also duplicated (Funabiki *et al*., 2023) and the absence of one *CDCA7* paralog in mice had no effect on TE silencing (Vukic *et al*., 2024), suggesting that *CDCA7* paralogs also have redundant functions in animals. In *A. thaliana*, *CDCA7α* has a stronger impact on mCG than *CDCA7β*. This difference is unlikely due to the divergence in their C-terminal domains, as this region was found to be dispensable for TE silencing. Instead, the stronger impact of *CDCA7α* likely results from its higher expression levels. The redundant roles of *CDCA7α* and *CDCA7β* indicate that *CDCA7* duplication may have increased CDCA7 protein dosage or enhanced robustness against genetic and environmental challenges (Kuzmin, Taylor and Boone, 2022).

How does CDCA7 promote mCG? Studies in *Xenopus* egg extracts and mouse embryonic stem cells showed that CDCA7 recruits HELLS (the DDM1 ortholog) to chromatin and activates its nucleosome remodeling activity (Jenness *et al*., 2018; Hardikar *et al*., 2020; Shinkai *et al*., 2024; Wassing *et al*., 2024). Moreover, the zinc finger of animal CDCA7 and *A. thaliana* CDCA7α bind hemi-methylated CG dinucleotides, which are formed during DNA replication (Hardikar *et al*., 2024; Shinkai *et al*., 2024; Wassing *et al*., 2024). Therefore, CDCA7 likely functions by recruiting DDM1/HELLS to hemi-methylated mCG sites (Wassing *et al*., 2024) and DDM1/HELLS would then remodel nucleosomes to provide methyltransferases with access to DNA (Zemach *et al*., 2013; Lyons and Zilberman, 2017). This model is compatible with our observations, implying that this CDCA7 function is conserved in plants and metazoans.

CDCA7α/β act as facultative activators of DDM1, since they act through DDM1, but their absence had a weaker impact on mCG than DDM1 loss. This suggests that DDM1 may function on its own or with other factors than CDCA7α/β. DDM1 alone can remodel nucleosomes *in vitro* (Brzeski and Jerzmanowski, 2003; Lee *et al*., 2023; Osakabe *et al*., 2024), but it is not known whether DDM1’s intrinsic activity is sufficient *in vivo* or whether protein partners like CDCA7α/β are required to stimulate nucleosome remodeling. In contrast, HELLS requires CDCA7 for *in vitro* activity (Jenness *et al*., 2018; Nartey, Goodarzi and Williams, 2023) and animals have limited amounts of non-CG methylation (Ziller *et al*., 2011). This distinction suggests that DDM1’s intrinsic activity may have evolved to maintain non-CG methylation independently of CDCA7α/β, which fits with our observation that CDCA7α/β had a specific effect on mCG.

Beyond mCG maintenance, DDM1/HELLS regulates histone variant exchange in plants and animals (Ni *et al*., 2020; Ni and Muegge, 2021; Osakabe *et al*., 2021; Jamge *et al*., 2023; Lee *et al*., 2023), but the contribution of CDCA7 to this function had not been previously investigated. Our results indicate that CDCA7α/β act specifically on mCG regulation, with minimal impact on other DDM1 functions. This specificity aligns with the fact that *CDCA7* has been lost in most eukaryotes lacking DNA methylation, while *DDM1/HELLS* has been retained (Funabiki *et al*., 2023). Accordingly, the absence of one CDCA7 paralog affects mCG but not H3K9me3 in mice (Vukic *et al*., 2024). Therefore, the mild reduction of heterochromatic marks other than mCG in *cdca7α/β* mutants is likely an indirect effect of decreased mCG. This raises the question of how CDCA7α/β selectively contributes to a single DDM1 function, given that DDM1-dependent heterochromatin marks generally co-exist in the genome. We propose a model in which histone variant incorporation by DDM1 occurs independently of CDCA7α/β after DNA synthesis. Subsequently, CDCA7α/β recruit DDM1 at hemi-mCG sites to stimulate nucleosome remodeling and allow mCG deposition. This model aligns with kinetics of nucleosome assembly being faster than mCG deposition after DNA replication (Stewart-Morgan, Petryk and Groth, 2020; Stewart-Morgan *et al*., 2023), and is further supported by observations that CDCA7 preferentially binds hemi-mCG sites located in nucleosomal rather than linker DNA (Wassing *et al*., 2024). Further studies will be essential to pinpoint the specific timing and coordination of CDCA7 in DDM1/HELLS functions.

We provide evidence that a *trans*-factor, *CDCA7α*, causes changes in DNA methylation that in turn, have the capacity to influence natural populations. Changes in expression levels of this limiting factor perturbed mCG levels on TEs and potentially affected seed size, an adaptive trait. Our results support the hypothesis that evolutionary changes in the dosage of epigenetic regulators have caused global changes in epigenetic profiles(Baduel and Sasaki, 2023). To date, we have identified several other *trans*-regulators controlling mCH, including *CMT2*, *CMT3*, *MIRNA823* (which targets *CMT3* transcripts), *JMJ26*, RNA polymerase V (Dubin et al. 2015; Kawakatsu et al. 2016; Sasaki et al. 2019; Sasaki et al. 2022). To this list, we now add *CDCA7α* and *DDM1*, affecting CG, which are directly inherited. As combinations of epigenetic mutants often exhibit epistasis (Cao and Jacobsen 2002; He et al. 2022; Shimada et al. 2024), these allelic interactions—spanning multiple pathways and all DNA methylation contexts—considerably elevate epigenetic diversity and have the potential to modify gene regulatory networks and TE silencing. Such genetically inherited epigenetic variation can become a target of natural selection (Baduel and Sasaki, 2023). Our study has provided insights into epigenome variation historically observed in nature at the molecular level. Further studies, including analysis of genetic and epigenetic architecture in other species, will enhance our understanding of the complex ecological significance of the epigenome in the tree of life.

## Materials and Methods

### Plant material

Seeds sown on soil were stratified at 4°C for two days in the dark and grown at 21°C under long-day conditions (16 h light and 8 h dark) in a climatic chamber.

#### Mutants from stock centers and previous studies

*cdca7α-1* (SALK_100123C), *cdca7α-2* (SALKseq_124190.3), *ddm1-2* (Vongs *et al*., 1993) mutant and control plants were in the Col-0 genetic background. To reset the effect of the transgenerational aggravation of the *ddm1-2* mutant phenotype (Kakutani *et al*., 1996), we backcrossed a *ddm1-2* homozygous mutant six times to Col-0, keeping *ddm1-2* heterozygous throughout, and subsequently isolated homozygous *ddm1-2* mutants which were considered first generation for this study.

#### CRISPR-Cas9 mutagenesis

CRISPR mutants were generated by transforming Col-0, *ddm1-2* second-generation mutants, or *cdca7α-2* mutants. To mutate the *CDCA7α/β* genes, we used a CRISPR-Cas9 system based on an intronized *Cas9* gene controlled by the ubiquitous RPS5 promoter (Grützner *et al*., 2021). Guide RNAs (listed in Supplementary Table 2) controlled by a U6 promoter were cloned into pICH47751 or pICH47761, and binary plasmids were assembled using golden gate assembly with one or two guide RNAs, the intronized Cas9 gene, a linker for plasmid assembly, and a seed coat fluorescent reporter using mVenus (Bensmihen *et al*., 2004) or a Basta resistance gene. Transgenic lines were generated with *Agrobacterium tumefaciens* GV3101 by floral dipping. T1 transformed seeds were selected by Basta resistance or seed coat fluorescence. Mutations were identified by PCR and Sanger sequencing. We isolated T2 seeds devoid of the transgene and selected mutations. We obtained stable homozygous mutant lines in T2 or T3, and used their progeny for this study. Details about primers, guide RNAs and CRISPR mutations used in this study can be found in Supplementary Tables 2 and 3.

#### Complementation

*CDCA7α* and *CDCA7β* transgenes were cloned from genomic DNA to include endogenous regulatory sequences in promoters, 5’UTRs, and introns. We used the Greengate cloning system (Lampropoulos *et al*., 2013) to assemble constructs into a pGGSun backbone (Incarbone *et al*., 2023). Promoters included 5’ UTRs and were cloned from the first base before the ATG start codon to 792 bp upstream for *CDCA7α* and 1349 bp upstream for *CDCA7β*. The alternative *CDCA7α_a_* promoter was cloned from *A. thaliana* accession 6966 using the same primers as for *CDCA7α_ref_* promoter, resulting in an 831 bp fragment. Coding sequences contained all introns and exons, except the 3’ UTR and the start and stop codon, according to the tag position. Endogenous *CDCA7β* BsaI sites were removed by mutagenesis, introducing silent mutations (GTC to GTT and GGT to GGA). N-terminal tags had a start codon with 3xcMyc fused with mTurquoise2, C-terminal tags had mTurquoise2 fused with 3xcMyc and a stop codon. All constructs were cloned with the terminator of the UBQ10 gene (628 bp) and a mVenus seed coat reporter gene. Deletions and point mutations were introduced in constructs using in vivo cloning (García-Nafría et al. 2016). Transgenic lines were generated with *A. tumefaciens* GV3101 by floral dipping and transformed seeds were selected by seed coat fluorescence (Bensmihen *et al*., 2004).

### Phylogenetic Analysis of *CDCA7*

To reconstruct the evolutionary history of CDCA7, we constructed orthologous gene clusters (i.e., orthogroups) from two different proteome datasets using sequential, dense taxon sampling across two levels: Archaeplastida (50 species) and Viridiplantae (66 species, enriched for Brassicales) (Supplementary Table 4). This method enabled us to estimate more precisely the timing of the *CDCA7α/β* duplication event. Orthofinder v2.5.2 (Emms and Kelly, 2019) was used to cluster genes in a non-biased manner by comparing each gene to the entire proteome dataset, and DIAMOND (Buchfink, Reuter and Drost, 2021) for homology search (‘-S diamond_ultra_sens’). For the Archaeaplastida-level survey, we included previously-identified metazoan CDCA7s (Funabiki *et al*., 2023) as an outgroup. MAFFT v7.310 (Katoh and Standley, 2013) was used to align protein sequences based on orthogroups. Individual maximum likelihood gene trees were built with IQ-TREE v2.1.2 (Minh *et al*., 2020), which used model selection (‘-m MFP’) and an ultrafast bootstrap approximation approach (1,000 replicates). The phylogenetic trees were visualized using iToL v6.7 (Letunic and Bork, 2024).

We used ancestral sequence reconstruction to reconstruct the CDCA7 protein sequence prior to the Brassicaceae CDCA7α/β duplication event. Based on the phylogenetic positions, we selected 48 protein sequences from Brassicaceae CDCA7 homologs and Amborella CDCA7 as an outgroup and aligned them using MAFFT. Following the multiple sequence alignment, IQ-TREE was used to rebuild a phylogenetic tree that will be used as input for the initial ASR analysis. First, we converted the protein sequence alignment to an absence/presence matrix, then used IQ-TREE’s ancestral sequence reconstruction (’-asr’) option with the rooted phylogenetic tree. The absence/presence ancestral state output was used to trim residues from the protein sequence alignment based on the calculated posterior probability of each residue. The final ancestral sequence reconstruction used the trimmed alignment to generate the ancestral Brassicaceae CDCA7 protein sequence.

### Measuring flowering time and seed size

Flowering time was quantified by counting the number of mature rosette leaves at the anthesis of the first flower. Plants were grown in a randomized layout under short-day (8h light/16h dark) or long-day (16h light/8h dark) conditions at 21°C in a climatic chamber. The *cdca7α/β* mutants were in the 5th generation of inbreeding post-CRISPR transformation, while *ddm1-2* mutants were in the 3rd generation.

For seed size measurements, plants were grown in a randomized pattern from seed batches with reduced heterogeneity. Specifically, seeds were sorted on size to exclude the 5% of biggest and smallest seeds, using a Boxeed seed phenotyping robot (Labdeers). Four crosses were performed for each inflorescence, yielding 5 to 47 seeds per silique (mean = 31.08, n = 72). We used either the main floral stem, or the first axillary branch as a female inflorescence. A linear model showed that the mean seed surface area per silique was unaffected by the inflorescence type (P = 0.64) or the number of seeds per silique (P = 0.89), allowing comparisons to be made without accounting for these variables. Seed surface was measured using scanners and ImageJ scripts.

### Sample collection for epigenomics

Plants were grown following a randomized pot pattern. For methylomes and transcriptomes of *cdca7* and *ddm1 cdca7* mutants (**Fig. 2, 3**), we pooled rosette leaves from three 31-day-old plants (two rosette leaves each), ground the tissues in liquid nitrogen, and split the ground tissues for DNA and RNA extraction. For ChIP-seq, we pooled at least 5 plants using 5 rosette leaves each up to 1.5 g per sample. For methylomes and transcriptomes of complemented *cdca7* mutants (**Fig. 4, 5**), we used 28-day-old plants: one rosette leaf (2-3 cm long) was flash frozen first for RNA extraction, then all remaining rosette leaves except cotyledons and leaves 1 and 2 were collected for DNA extraction. All generated libraries are listed in Supplementary Table 5, with the number of inbreeding generations for each mutant.

### Whole genome bisulfite sequencing (WGBS)

#### Library construction and sequencing

To measure DNA methylation levels in *cdca7α-2, smp2,* and *AT4G37130* mutants (**Fig. 1e**), genomic DNA was extracted from rosette leaves at 9-true-leaf stage using the GeneJET Plant Genomic DNA Purification Kit (Thermo Scientific) and sheared with an E220 Focused-ultrasonicator (Covaris) to achieve an average fragment size of approximately 350 bp. Sequencing libraries were prepared using the NEBNext Ultra II DNA Library Prep Kit (New England BioLabs) with methylated adapters (New England BioLabs). The adapter-ligated DNA underwent bisulfite conversion using the EZ-96 DNA Methylation-Gold MagPrep Kit (Zymo Research). Bisulfite-treated samples were amplified using EpiMark Hot Start Taq DNA Polymerase and indexed with NEBNext Multiplex Oligos for Illumina (New England BioLabs). All libraries were sequenced on either an Illumina NextSeq 550 or HiSeq 2500 platform.

#### Estimation of DNA methylation levels

All reads were mapped on TAIR10 reference genome using a Methylpy pipeline v1.2 (https://github.com/yupenghe/methylpy). DNA methylation levels were estimated as weighted methylation levels for each transposon defined in Araport11 annotation. CMT2- and RdDM-targeted transposons were defined as having differential levels of methylation (> 0.1) between wild-type and *cmt2* or *drm1drm2* in Col-0 as previously described (Kawakatsu *et al*., 2016). For each line, average DNA methylation was calculated using all transposons for which at least one read was mapped.

### Tagmentation-based WGBS (T-WGBS)

#### Library construction and sequencing

Genomic DNA was extracted from rosette leaves with the DNeasy Plant Pro kit (Qiagen, reference 69206), following the manufacturer’s instructions, using 1 to 5 biological replicates per genotype (Supplementary Table 5). Illumina libraries of sodium bisulfite-converted DNA were prepared with a tagmentation-based approach (Wang et al. 2013) with modifications (Montgomery and Berger, 2023, bioRxiv). Briefly, we tagmented 50 ng of DNA with Tn5 (Molecular Biology Services, IMP, Vienna, Austria). Tagmented DNA was purified with the Zymo DNA Clean and Concentrator kit (Zymo Research) instead of SPRI beads. After oligonucleotide replacement, gap repair, and purification, we used the entirety of the eluted DNA for bisulfite conversion, which was performed with the EZ DNA Methylation-Gold Kit from Zymo (reference D5006). The rest of the protocol was performed as indicated by Wang et al (2013). Directional libraries were amplified with Nextera-compatible unique dual indices and sequenced on a NovaSeq 6000 instrument using an S4 flow cell to generate paired-end 150 bp reads with around 25 million reads per sample.

#### T-WGBS analysis

Libraries for *cdca7α/β* and *ddm1* mutants (**Fig. 2, 3**) were randomly subsampled to 30 million reads to improve depth homogeneity, since there were up to 2-fold more reads in some samples. The data were analyzed with the nfcore methylseq pipeline v2.3.0 (https://nf-co.re/methylseq) (Ewels *et al*., 2020) with Nextflow (v22.10.7). Cytosine positions covered by fewer than 3 reads (--meth_cutoff 3) were excluded. Tagmentation creates gaps in the double-stranded DNA which are later filled by unmethylated cytosines during gap repair. These nucleotides were excluded from the analysis by trimming reads with the "clip" and "ignore" options, using M-bias plots to ensure the removal of unmethylated fragment ends. The parameters were as follow: --meth_cutoff 4 --comprehensive --cytosine_report --clip_r1 14 --three_prime_clip_r1 4 --three_prime_clip_r2 2 --ignore_3prime_r2 17. Note that the unmethylated cytosines are incorporated at the 5’ end of read 2, while the options mentioned above suggest we ignored the 3’ end of read 2 instead, but methylseq v2.3.0 had an issue which made it clip the wrong end of read 2 (see https://github.com/nf-core/methylseq/issues/299). Using M-bias plots, we noticed a 5’ bias on read 1 that resembled the Tn5 sequence bias (Wolpe, Martins and Guertin, 2023), and removed it by clipping 14 nucleotides at the 5’ end of read 1. We also excluded additional small methylation biases at the 3’ end of read 1 and read 2.

To average biological replicates, we computed the mean methylation level for individual cytosine positions, including all replicates that met the minimum coverage threshold. Positions covered in only one replicate were retained to maximize data inclusion. To compute the average methylation values over annotation sets or 100 bp bins, we used the *bigWigAverageOverBed* script from UCSC tools (https://github.com/ucscGenomeBrowser/kent-core/tree/master). Only TEs covered in all samples from a common batch were retained. To calculate average values in bins (metaplots), annotations were scaled to an arbitrary size of 1000 bp, and the methylation value was calculated across annotation length by 25 bp bins, and up to 500 bp upstream and downstream. This was done with the computeMatrix scale-regions --binSize 25 and plotProfile functions from deepTools v3.3.1. To analyze 5mC changes in 100 bp bins, we segmented the genome in non-overlapping 100 bp bins, counted the number of each methylation context per bin using a custom python script. Average 5mC per bins was calculated with *bigWigAverageOverBed*, and only bins that were covered in all samples were retained for further analysis. To compute Spearman correlations, we kept bins which had at least 10% mCG, 6% mCHG and 5% mCHH in WT. Gene body methylation analysis was restricted to protein-coding genes covered in all samples which had more than 10% mCG, less than 10% mCHG and 10% mCHH in WT (n = 9938).

To analyze DNA methylation at TEs, we excluded TEs unmethylated in the WT, using a threshold of 5% methylation for mCG, 3% for mCHG and 2% for mCHH, after which sample size were of 18052, 16739 and 18037, respectively.

### 3’ tag-sequencing of messenger RNAs

#### Library construction and sequencing

Total RNA was extracted using the RNA isolation kit provided by the VBC core facilities. The kit uses a lysis step based on guanidine thiocyanate (adapted from (Boom *et al*., 1990) and carboxylate-modified Sera-Mag Speed beads and was applied using the KingFisher instrument (Thermo).

Libraries were prepared in 96-well plates using a custom-protocol described previously (Clavel *et al*., 2024). The oligo d(T) primers used for reverse transcription contain 8-bp unique molecular identifiers (UMIs), 7-bp barcodes for multiplexing, and a Truseq adapter. After reverse transcription, samples are pooled and RNA-cDNA duplexes tagmented with Tn5. Using primers matching the Truseq and Tn5 adapters, the 3’ ends of mRNA transcripts are subsequently amplified by PCR, while P5 and P7 sequences and plate-specific dual indices are added for Illumina sequencing.

We started from 80 ng of total RNA in 4 µl, adding 1 µl of 0.2 µM primers for reverse transcription. RNAs were denatured 3 min at 72°C and kept on ice. We mixed 1 µl of the SMARTScribe reverse transcriptase (Takara) in a final volume of 10 µl with 1x SMARTScribe buffer, 1.75 mM dNTPs, 2 mM DTT, 3 mM MnCl_2_ for 1h at 42°C followed by inactivation for 15 min at 70°C. All samples were pooled together and purified using 1.2x volume of SPRI beads (Molecular Biology Services, IMP, Vienna, Austria). After mixing and 2 min at room temperature, beads were collected on a magnet, washed two times with 70% ethanol, dried, and eluted in 60 µl of 10 mM Tris-HCl pH 8.0. To assemble transposomes, we mixed Tn5 (0.38 mg / ml) with 18 µM of annealed adapters, 50% glycerol and 1x Tn5 loading buffer (6x buffer: 300 mM Hepes KOH pH 7.2, 600 mM NaCl, 0.5 mM EDTA, 6 mM DTT, 0.6% Triton-X-100) and incubated for 1h at 37°C. For tagmentation, 16 µl of pooled RNA-cDNA was mixed with 16 µl of transposome and 32 µl of 2x tagmentation buffer (20 mM Tris (hydroxymethyl)aminomethane, 10mM MgCl2, adjusted to pH 7.6 and 20% (vol/vol) dimethylformamide). After 8 min at 55°C, we added 16 µl of 0.2% SDS and incubated 5 min at room temperature. Nucleic acids were purified with 2x SPRI bead volume as indicated above with a 48 µl elution. For PCR, we pre-incubated at 95°C the KAPA HiFi HotStart 2X ReadyMix for 90 s to activate the polymerase and amplified libraries in 100 µl final volume using 1 µM of primers and 46 µl of purified cDNA-RNA duplexes with the following program: 72°C for 3 min, 95°C for 30 sec, (95°C for 20 sec, 65°C for 30 sec, 72°C for 30 sec) for 12 cycles, 72°C for 5 min. Libraries were purified with 0.8x volume of SPRI beads as indicated above and eluted in 62 µl. After controlling for appropriate fragment size distribution on a 5200 Fragment Analyzer System (Agilent), libraries were sequenced as paired-end 150 bp reads on a NovaSeq S4 (**Fig. 2, 3**) or a NovaseqX 10B (**Fig. 4, 5**) flow cell.

#### Mapping and differential expression

Reads were demultiplexed, trimmed, collapsed to remove duplicates, mapped, and quantified. The code and documentation are available at https://github.com/pierre-bourguet/3-tag-rna-seq, but we provide the most important details below. Read1, which starts directly in the transcript at the site of Tn5 cutting, was used for mapping. Read2, which contains the sample-specific index, UMI, polyA tail, and eventually the 3’ end of the transcript, was used for demultiplexing and UMI extraction. Specifically, we used the first 7 bases of read2 to identify indices specific to each well in the plate, and the next 8 bases to extract UMIs. For read1, the 3’ end was trimmed for polyA and adapter contamination and reads shorter than 50bp were discarded. UMIs were prepended to read1 and these chimeric reads were used to collapse duplicates into a single read with clumpify (bbmap v38.26) (Bushnell, 2014), with no mismatches allowed in the UMI and two mismatches allowed in the transcript sequence. Mapping was performed with STAR v2.7.11a (Dobin *et al*., 2013) and mapped reads were counted with Salmon v1.10.1 in alignment mode (Patro *et al*., 2017) in sense and antisense. We used the TAIR10 genome and AtRTD3 transcriptome reference (Zhang *et al*., 2022). Because the AtRTD3 GFF and fasta files were sometimes discordant, we regenerated a fasta file from a fixed GFF file using AGAT v1.0.0. We observed that the 3’-end of transcripts were often located outside of transposable element gene annotations, likely because they do not include 3’ UTRs, but within TAIR10 TE annotations. For this reason, and to avoid overlapping annotations, we removed “transposable element gene” annotations from AtRTD3 and added the 31189 transposable element annotations from TAIR10 instead.

We used DESeq2 v1.42.1 (Love, Huber and Anders, 2014) for a quantitative analysis of read counts. For a given feature, counts in the sense or antisense orientation were considered as independent features. We ran the DESeq2 model in single-strand mode, removing features which did not have at least 10 reads in a minimum of 3 samples. Features were considered differentially expressed at a FDR of 10% (adjusted p-value ≤ 0.1) and an absolute log_2_ fold change relative to the WT equal or greater than 1. To avoid confounding effects of overlapping annotations, we removed TEs misregulated in sense and antisense if they overlap (1 bp or more) a protein-coding gene in the same or reverse orientation, respectively. We use DESeq2 to normalize counts with the median ratio method and apply a variance-stabilizing transformation (VST).

To quantify transgene expression, the transgene sequence and annotations were added as an additional chromosome in the fasta and GTF files. The C-terminal mTurq2:3xcMyc tag allowed us to distinguish transgenic reads from endogenous *CDCA7α* reads. For Fig. 5e, a two-way ANOVA showed a significant effect of the promoter haplotype (*p-*value = 5e-6) on transgene expression levels, but no effect of the genetic background (*p-*value = 0.46) and no interaction (*p-value* = 0.48) and therefore, we merged the data from the two different backgrounds.

### Chromatin Immunoprecipitation followed by sequencing

#### Library construction and sequencing

1.5 g of rosette leaves from 31-d-old plants were fixed in 37 ml 1% formaldehyde in PBS by vacuum-infiltration for 10 min. The cross-linking reaction was quenched by adding glycine to a final concentration of 125 mM. Tissues were frozen in liquid nitrogen and ground in a 5-ml grinding jar on a Retsch Mixer Mill MM 200 at 30 Hz for 45 s. Tubes were kept at room temperature until the powder showed signs of thawing. We added 35 ml of nuclei isolation buffer (NIB, 10 mM MES-KOH pH 5.3, 250 mM sucrose, 10 mM NaCl, 10 mM KCl, 2.5 mM EDTA, 0.1 mM spermine, 0.1 mM spermidine, 2.5 mM ß-mercaptoethanol, 0.3% Triton X-100 and protease inhibitors) (Lorkovic 2004 MBC), followed by vortexing until homogeneous. The extract was filtered through two layers of Miracloth, which were washed with 10 ml of NIB and spun at 3,400 rpm at 4°C for 5 min. Pellets were resuspended in 25 ml of NIB, vortexed and incubated for 10 min on ice to fully dissolve chlorophyll content. Tubes were spun as above and pellets were carefully washed with 5 ml of shearing buffer (10 mM Tris-HCl pH 8.0, 1 mM EDTA, 0.1% SDS, and protease inhibitors) without resuspending and spun again. This washing step was repeated and pellets were resuspended in 0.9 ml of shearing buffer. One ml of the extracted chromatin was transferred to Covaris glass tubes and sonication was conducted at 4°C for 900 seconds per sample (peak power 140.0; duty factor: 5.0; Cycles/Burst: 200). Sheared chromatin was spun for 5 min at 4°C at 15,000 rpm. The supernatant was collected and spun again, and the supernatant was transferred to a 5 ml tube. Fifteen µl of chromatin were kept to control for sonication efficiency.

Chromatin was diluted to 3.5 ml to reduce SDS concentration with a ChIP dilution buffer (16.7 mM Tris-HCl pH 8.0, 167 mM NaCl, 1.2 mM EDTA, 1.1% Triton X-100, 0.01% SDS and protease inhibitors). For preclearing, 200 µl of Dynabeads protein A (Thermofisher, reference 1001D) were added to the chromatin and rotated 1 h at 4°C. Beads were discarded by collecting supernatants after two centrifugations at maximum speed for 30 s. A hundred µl of pre-cleared chromatin was kept at -20°C for input control, and the remaining chromatin was aliquoted with 500 µl per immunoprecipitation (IP) and incubated with 5 µg of anti-H3 (Abcam, ab1791), anti-H2A.W.6/7, anti-H2A.Z.9/11, anti-H3K27me1 (Millipore, 17-643) or anti-H3K9me2 (Abcam, ab1220) antibodies at 4 °C overnight with rotation. After incubation, samples were mixed with 30 µl of protein A magnetic beads, rotated at 4°C for 3 h, and washed two times with a low-salt buffer (20 mM Tris-HCl pH 8.0, 150 mM NaCl, 2 mM EDTA, 1% Triton X-100 and 0.1% SDS), once with a high-salt buffer (20 mM Tris-HCl pH 8.0, 500 mM NaCl, 2 mM EDTA, 1% Triton X-100 and 0.1% SDS), once with a LiCl buffer (10 mM Tris-HCl pH 8.0, 1 mM EDTA, 0.25 M LiCl, 1% IGEPAL CA-630 and 0.1% deoxycholic acid) and twice with TE buffer (10 mM Tris-HCl pH 8.0 and 1 mM EDTA). Elution was done with 200 µl of 0.1 M NaHCO3 and 1% SDS, incubated at 65°C for 15 min. We added 20.4 µl of reverse cross-link buffer (0.4 M Tris-HCl pH 8.0, 2 M NaCl, 10 mM EDTA, 0.6 mg ml−1 proteinase K (Thermo Fisher Scientific)) and incubated at 45°C for 3 h and 65°C for 16 h. RNA was subsequently degraded for 30 min at room temperature with 10 µg of RNase A (Thermo Fisher Scientific), and DNA was purified with the ChIP DNA Clean & Concentrator kit (Zymo, reference D5205).

Libraries were prepared with the NEBNext Ultra II DNA library prep kit for Illumina (New England Biolabs), following the manufacturer’s instructions. Fragment size distribution was analyzed on a 5200 Fragment Analyzer System (Agilent), and size selection with SPRI beads (Molecular Biology Services, IMP, Vienna, Austria) was applied to remove large fragments and self-ligated adapters, when necessary. To maximize comparability, if only one sample had undesired fragments, we size-selected all samples for a given antibody. For size selection, briefly, DNA was diluted to 100 µl, we mixed in 60 µl of magnetic SPRI beads, collected the supernatant, and added 40 µl of magnetic SPRI beads, washed two times with 80% ethanol, and eluted DNA. Sequencing was carried out on a NovaSeq 6000 instrument using an S4 flow cell to generate paired-end 150 bp reads with around 25 million reads per sample.

#### ChIP-seq analysis

The data were analyzed using the nfcore/chipseq pipeline v2.0.0 (https://nf-co.re/chipseq/2.0.0/) (Ewels *et al*., 2020) with Nextflow (v22.10.7). We used the TAIR10 genome, custom arguments for read length (150 bp), fragment size (275), and blacklisted mitochondrial and chloroplast genomes. To get the average value for each annotation, we used the bigWigAverageOverBed script from UCSC tools (https://github.com/ucscGenomeBrowser/kent-core/tree/master). We further analyzed the data with deepTools v3.3.1 (Ramírez *et al*., 2016). To normalize by H3 or by the WT, we used *bigwigCompare* with a bin size of 10 bp, ignoring non-covered regions with *skipNonCoveredRegions*. To produce average profile plots (metaplots), annotations were scaled to an arbitrary size of 1000 bp and the enrichment value was calculated by 50 bp bins, using *computeMatrix scale-regions*.

### RT-qPCR

We used 21-d-old and 27-d-old rosette leaves to perform experiments in **Fig. S5d** and **Fig. S7b**, respectively. RNA extraction, transcript quantification, and normalization were performed as previously reported (Bourguet *et al*., 2020), except that reactions were run on a Roche^TM^ LightCycler® 96 system with the following program: 10 min at 45°C, 2 min at 95°C, and 40 cycles of 5 s at 95°C, 10 s at 60°C, and 5 s at 72°C. Primers are listed in Supplementary Table 2.

### Protein structure prediction

We used Alphafold3 (Abramson *et al*., 2024) to model protein structures and interactions, with the random seed set to 1. The top-ranking prediction was selected for representation. The predicted structure of CDCA7α with three Zn^2+^ ions was superimposed to *H. sapiens* CDCA7 in a complex with non-B-form DNA containing 5mC (PDB ID 8TLK) (Hardikar *et al*., 2024). Structures of DDM1 with a nucleosome (**Fig. S8**) were predicted with the H2A.W.6 and H3.1 histone variants, using the exact sequences from a cryo-EM structure of DDM1 (PDB ID 8J90) (Osakabe *et al*., 2024). Precisely, protein chains were from the HTA6, HTR2, AT2G28740 (H4) and HTB9 proteins, with a Widom 601 DNA sequence. The best ranked prediction was in good agreement with the 8J90 structure, as root mean square deviation (RMSD) between 190 pruned atom pairs was 1.043 angstroms. Unstructured regions were removed from the visualization for clarity. To find interacting residues between CDCA7α/β and DDM1 (**Fig. 4d, g**, **S8d**), we used AlphaFold-Multimer (Evans *et al*., 2022) using a local implementation of Colabfold (Mirdita *et al*., 2022) and the top-ranking prediction. Graphical representations and analyses of structures were done with ChimeraX (Meng *et al*., 2023).

### Recombinant protein expression

#### Cloning

CDCA7α and CDCA7β cDNAs were amplified by RT-PCR with primers indicated in Supplementary Table 2, using RNA purified from WT flowers. We used the canonical CDCA7α.2 transcript variant according to AtRTD3 (Zhang et al. 2022). cDNAs were cloned into pGEX-4T-1 (Cytiva) using *EcoRI*/*SalI*. DDM1 cDNA into pET15b was previously described (Osakabe et al. 2021).

#### Expression and purification of recombinant proteins and *in vitro* pull-down experiments

BL21 (DE3) RIL *E. coli* cells transformed with plasmids for protein expression were grown at 37°C overnight in 100 ml LB. Cultures were diluted in 1 L of LB and grown for three hours at 20°C and then induced for 7 hours at 20°C with 1 mM IPTG. For GST-tagged proteins cell pellets were resuspended in 20 ml of extraction buffer (50 mM Tris-HCl pH 8.0, 1 M NaCl, 1 mM DTT, 0.1% Triton X-100) containing protease inhibitors (Roche), 10 µl of benzonase (1 mg/ml) and 50 mg of lysozyme. After sonication (Bioruptor, Diagenode) for 10 min at high intensity (5’’ on / 5’’ off) and 5 min at medium intensity (5’’ on / 5’’ off), extracts were centrifuged for 15 min at 4°C at 40,000 × *g*. Extracts were incubated with 500 µl of glutathione Sepharose 4 fast flow (Cytiva) at RT for one hour and then transferred to disposable columns and washed with 5 column volumes of extraction buffer. Proteins were eluted with six 300 µl elution steps with 50 mM Tris-HCl pH 8.0, 500 mM NaCl buffer containing 20 mM reduced glutathione and 1 mM DTT.

For purification of His_6_- or His_6_SUMO-tagged DDM1, cell pellets from 2 L cultures were resuspended in 25 ml of extraction buffer (50 mM Tris-HCl pH 7.5, 500 mM NaCl, 2 mM DTT, 0.05% NP-40) containing protease inhibitors (Roche), 10 µl of benzonase (1 mg/ml) and 50 mg of lysozyme and processed as for GST-tagged proteins. Extracts were incubated with 500 µl of Ni-NTA beads (Qiagen) for one hour at RT and then transferred to disposable columns and washed with 5 column volumes of extraction buffer containing 5 mM imidazole but without benzonase and lysozyme. Proteins were eluted with six 400 µl elution steps with elution buffer (50 mM Tris-HCl pH 7.5, 500 mM NaCl, 2 mM DTT, 300 mM imidazole).

In all purifications, elution fractions were analyzed on 10-12% SDS-PAGE, pooled, and buffer was exchanged into 50 mM Tris-HCl pH 7.5, 100 mM NaCl, 1 mM DTT by centrifugation over Amicon Ultra-15 30 kDa and 50 kDa cut-off centrifugal filters (Millipore) for GST-tagged CDCA7 and His-tagged DDM1, respectively.

For pull-down, equimolar amounts of GST-tagged proteins along with GST alone were mixed with 5 µg of His_6_SUMO-DDM1 and incubated with 10 µl of magnetic glutathione beads (Thermo Fisher Scientific) for 90 min at RT in binding buffer (20 mM Tris-HCl, pH 7.5, 100 mM NaCl, 1 mM DTT, 0.1% Triton X-100). Beads were washed 6 times with binding buffer, denatured in 30 µl of 1 × SDS-PAGE loading buffer and 10 µl were loaded on 10% SDS-PAGE and analyzed by western blotting with anti-DDM1 antibody. Input lanes were loaded with 1/20 of protein used for pull-down.

### Genome-wide association studies

GWAS were conducted using LIMIX (Lippert *et al*., 2014) version 3.0.4 with full genome SNPs from the 1001 genome project (10,709,949 SNPs for mCG levels and 3,686,507 SNPs for gene expression) and the following linear mixed model (LMM),

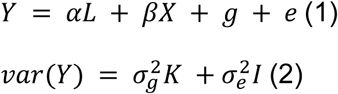

where *Y* is the vector of a phenotype, fixed terms, *L* and *X*, are *n* × 1 vectors corresponding to a cofactor and a genotype to be tested (SNP) with the parameters *α* and *β*, respectively. 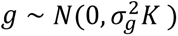 and 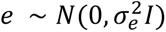 are random terms, including the identity-by-state (IBS) kinship *K* matrix representing the genetic relatedness (Yu *et al*., 2006; Kang *et al*., 2008) and the residuals, respectively. The vectors *Y*, *L*, and *X* were z-scored. Models without a cofactor take *α* = 0. SNPs with MAF ≥ 5% were used for association studies. Bonferroni correction was applied for multiple-testing correction after excluding all SNPs with MAF < 5%.

### Mediation analysis and heritability

The fraction of genetic effects on mCG variation mediated by *CDCA7α* expression was estimated using mediation analysis with the published program (Sasaki, Frommlet and Nordborg, 2018). Two alleles, *CDCA7α_a_* and *CDCA7α_b_*, were fitted into one model together as a single genotype vector: *CDCA7α*_Ref_ :0, *CDCA7α_a_* :1, and *CDCA7α_b_ :2*.

### Local genetic structure

#### Genome alignment of *CDCA7α* region and haplotype network construction

Fully assembled genome sequences of 27 *A. thaliana* lines (Igolkina *et al*., 2024), *A. lyrata* v1.0, *A. halleri* v2.1.0, and *Capsella rubella* v1.0 (Slotte *et al*., 2013) were used for the alignments of diverged *CDCA7α* promoter region. *A. lyrata* and *C. rubella* genomes were downloaded from NCBI, and *A. halleri* genome was downloaded from Phytozome (https://phytozome-next.jgi.doe.gov/). We extracted a 4 kb region covering the *CDCA7α* and *SMP2* coding sequences based on homology (chr4: 17,484,000–17,488,000 in the *A. thaliana* Col-0 genome, TAIR10) and made the alignment using CLC Sequence Viewer 8 with a gap open cost of 10.0 and a gap extension cost of 1.0. After removing all gaps, the alignment was applied to haplotype network construction by minimum spanning network (Paradis, 2018) implemented in popart (Leigh and Bryant, 2015).

The length of the intergenic region between *CDCA7α* and *SMP2* was estimated as the distance of one bp outside the start codons of *CDCA7α.1* and *SMP2.1* (chr4: 17,486,198–17,486,678; 481 bp in Col-0).

#### Local genetic structure of *DDM1*

The local genetic structure around the DDM1 locus was analyzed using principal component analysis (PCA) according to a previous study (Sasaki *et al*., 2021). Using the prcomp() function in R, we analyzed 475 SNPs from 1,135 lines in the 1001 Genomes Project (Kawakatsu et al. 2016), covering the region from chr5:26649000 to 26658200. Clusters represented by PC1 and PC2 were selected as components indicating the major haplotypes.

### Population structure and geographic distribution

We used the population structure information as genetic groups for 1135 lines from the 1001 genome dataset. The information for the additional 188 lines, including Tanzania and the Yangtze river populations, was extracted from a previous study (Hsu, Lo and Lee, 2019). The pairwise genetic distance matrix calculated by Hsu et al. (2019) was used to construct the neighbor-joining tree.

Latitude and longitude coordinates for each line were collected from the previous studies (1001 Genomes Consortium, 2016; Zou *et al*., 2017), and altitude information was estimated using the R package elevatr (https://github.com/USEPA/elevatr).

Local maps (Northern Sweden, The Iberian peninsula, and Yangtze Liver) and the elevation profile view were generated using QGIS 3.38.2 (https://www.qgis.org/).

### Public data analysis

#### Molecular phenotypes in natural populations

DNA methylation data used in this study was previously described (Kawakatsu *et al*., 2016; Sasaki *et al*., 2019). Briefly, bisulfite sequencing data for 774 lines in leaf tissue grown under ambient temperature was used for this analysis. Reads were mapped on each pseudogenome, using a Methylpy pipeline (https://github.com/yupenghe/methylpy). mCG levels were calculated as weighted methylation levels (Schmitz and Ecker, 2012) for individual TEs. We used TAIR10 annotation to define TE regions and 6,378 TEs having mapped reads in the region in all lines were used for all analyses. CMT2 and RdDM-targeted TEs were defined based on differential methylation level (>0.1) between wild-type and *drm1drm2* or *cmt2* in Col-0 (Stroud *et al*., 2013).

Expression data for 461 lines, collected in the 1001 Genomes Project under the same conditions as the DNA methylation data, were used in this study (Kawakatsu *et al*., 2016; Kornienko *et al*., 2023). As described in (Kornienko *et al*., 2023), raw RNA-seq data were aligned to the TAIR10 genome using STAR v.2.9.6 (Dobin et al. 2013) and gene expression levels were quantified using featurecounts (Liao et al. 2014).

#### Seed size phenotypes in natural populations

Seed size and altitude information in the Spanish local population were collected from (Vidigal *et al*., 2016). After extracting lines sequenced in the 1001 genome project, we analyzed only genetically Spanish group to exclude the global population structure.

#### *CDCA7α* and *CDCA7β* expression profiles

To analyze transcript levels of *CDCA7α* and *CDCA7β* across tissues and developmental stages (Hofmann, Schon and Nodine, 2019), we kept only tissues from the original dataset (Klepikova *et al*., 2016) and from (Hofmann, Schon and Nodine, 2019), to limit overrepresentation of embryonic samples, and filtered out senescent tissues.

### Data processing, plots and statistical analysis

All analyses were performed using R Statistical Software (R v4.3.2; https://www.R-project.org/) with the following packages: tidyverse 2.0.0, ggplot2 3.5.1 (Wickham, 2016; Wickham *et al*., 2019), patchwork 1.2.0, agricolae 1.3-7, ggbeeswarm 0.7.2, MASS 7.3-60 (https://patchwork.data-imaginist.com, https://cran.r-project.org/package=agricolae, https://cran.r-project.org/package=ggbeeswarm, https://cran.r-project.org/package=MASS). Tukey’s HSD tests were performed using one-way ANOVA with the aov() function, followed by the TukeyHSD() function. Heatmaps were built with the ComplexHeatmap package (Gu, Eils and Schlesner, 2016). Clustering was performed with k-means, using the consensus clustering from 1000 repeats.

## Data availability

The data described in this publication have been deposited in NCBI’s Gene Expression Omnibus and are accessible through GEO Series accession number GSE284119 (T-WGBS), GSE283989 (WGBS), GSE283987 (ChIP-seq), GSE284067 (3’ tag mRNA-seq).

## Acknowledgments

The authors express their gratitude to Akihisa Osakabe and Hironori Funabiki for the initial suggestion; Aleksandra E Kornienko and Almudena Molla Morales for providing DNA and RNA from *A. thaliana* accessions; Yoav Voichek for the 3’ tag-seq protocol; Vikas Shukla for the original tag-seq script; Marion Clavel, Nenad Grujic and Yasin Dagdas for GreenGate and CRISPR constructs; Tal Dahan-Meir and Thomas James Ellis for help with seed size measurements; Tatsuo Kanno for *ddm1* inbred lines; Jurah Alej for the local AlphaFold Multimer implementation; Elizaveta Grigoreva and Pierre Baduel for sharing genotype information of *A. lyrata* and *A. halleri*; Yoav Voichek, Olivier Mathieu, Leandro Quadrana for fruitful discussions; the Berger and Nordborg group for constructive feedback. For their support, we thank the staff at the Next Generation Sequencing and PlantS facilities at the Vienna BioCenter Core Facilities (VBCF), and the Molecular Biology Services from the Institute for Molecular Pathology (IMP). The computational results presented were obtained using the CLIP cluster (https://clip.science). This research was supported by FWF grants to F.B. P32054, P33380, PAT6138924; European Union’s Framework Programme for Research, Innovation Horizon 2020 (2014–2020) under the Marie Curie Skłodowska Grant Agreement no. 847548 (VIP2) to P.B, and Japan Society for the Promotion of Science to E.S. (JSPS 20 K22671 and 21H02538).

## Authors contributions

P.B., Z.L, D.C.K. and V.B. performed the experiments. P.B., E.S., C.H.C, A.I., and C.R.L. analyzed the data. P.B., E.S., F.B. and M.N. supervised the study and designed the experiments. P.B., E.S., F.B. and M.N. wrote the manuscript.

## Notes

### Competing Interest Statement

The authors have declared no competing interest.

