## Supplementary Figures S1 to S11. for "Major alleles of *CDCA7α* shape CG-methylation in *Arabidopsis thaliana*"

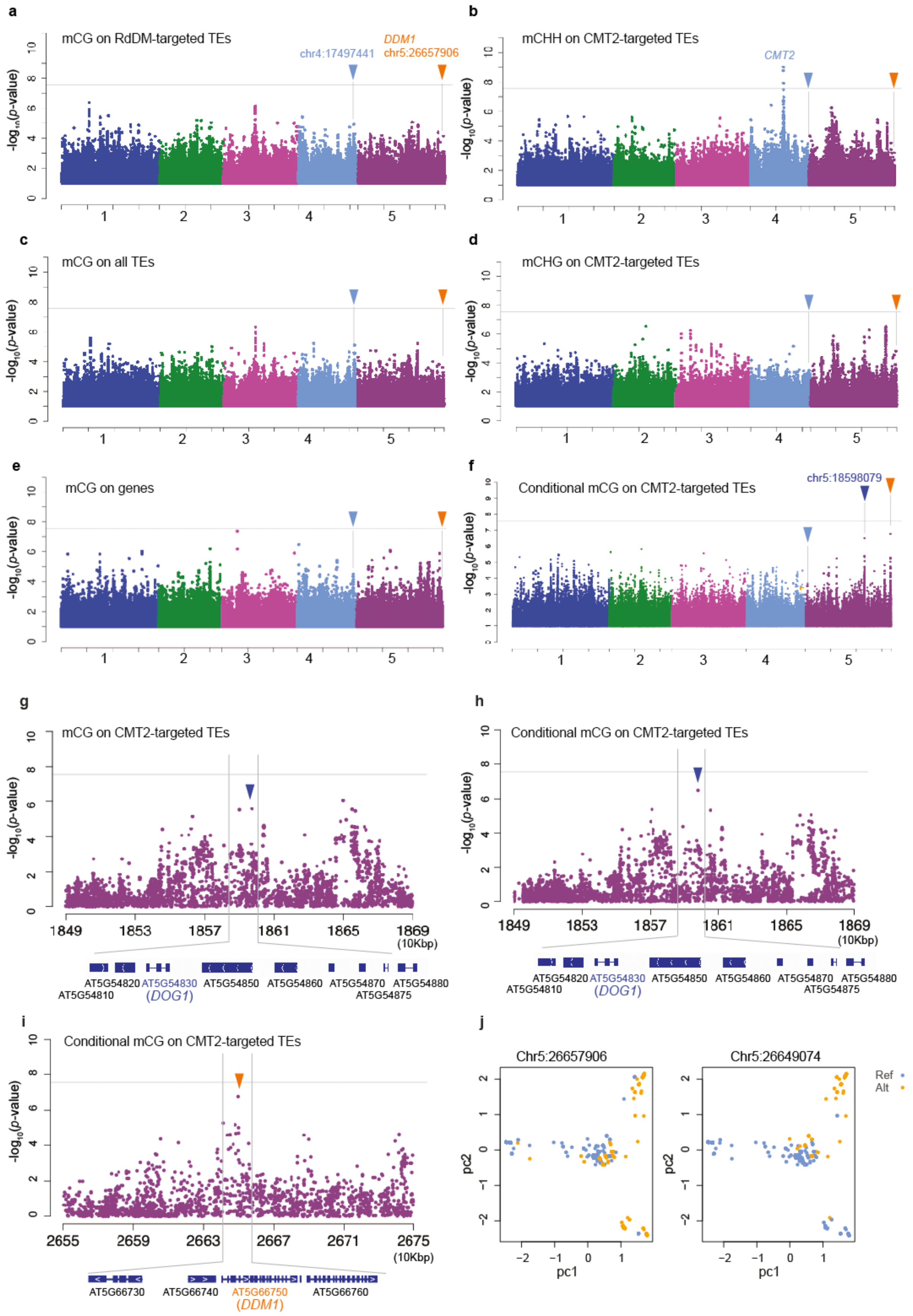
a
mCG on RdDM-targeted TEs
DDM1
chr4:17497441
chr5:26657906
b
mCHH on CMT2-targeted TEs
CMT2
c
mCG on all TEs
d
mCHG on CMT2-targeted TEs
e
mCG on genes
f
Conditional mCG on CMT2-targeted TEs
chr5:18598079
g
mCG on CMT2-targeted TEs
h
Conditional mCG on CMT2-targeted TEs
-log10(p-value)
1849 1853 1857 1861 1865 1869
(10Kbp)
AT5G54820 AT5G54830 AT5G54850 AT5G54860 AT5G54870 AT5G54880
AT5G54810 (DOG1) AT5G54875
i
Conditional mCG on CMT2-targeted TEs
2655 2659 2663 2667 2671 2675
(10Kbp)
AT5G66730 AT5G66740 AT5G66750 AT5G66760
(DDM1)
j
Chr5:26657906
Chr5:26649074
Ref
Alt
pc1
pc2

**Figure S1. The genetic basis of several genomic regions and various DNA methylation contexts.** GWAS for mCG on RdDM-targeted TEs (**a**), all TEs (**c**), genes (**e**), and CMT2-targeted TEs, conditioned on the genotype of chr4:1749744 (**f**). GWAS for mCHH (**b**) and mCHG (**d**) on CMT2-targeted TEs. Zoom-in figures compare a GWAS peak for mCG on CMT2-targeted TEs around chr5:18598079 (**g**) with the conditioned peak based on the genotype at chr4:1749744 (**h**). The conditioned peak at chr5:26657906 is shown in (**i**). Arrows on the Manhattan plots indicate SNP positions at chr4:17497441 (light blue), chr5:1859079 near *DOG1* (blue), and chr5:26657906 near *DDM1* (magenta), respectively. **b** and **d** are replotted from Kawakatsu et al. (2016), and **e** is replotted from Sasaki et al. (2022). j, local genetic structure of *DDM1* locus by PCA. Lines carrying reference and alternative alleles are mapped for chr5:26657906 (left) and chr:26649074 (right).

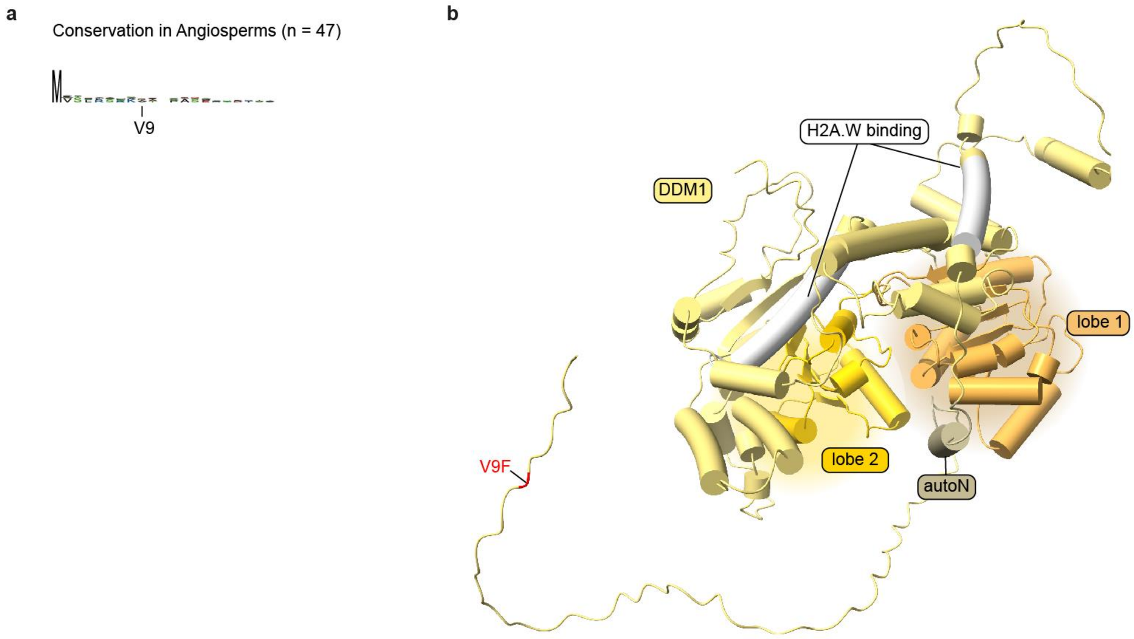


**Figure S2. Substitutions in the DDM1 alternative allele affect an aminoacid of unknown function** **a**, DDM1 consensus sequence among Angiosperms (n = 47), showing the position of the amino acid substituted in the *DDM1* alternative allele. **b**, Predicted DDM1 structure, highlighting the amino acid substituted in the *DDM1* alternative allele (red) and the different DDM1 domains. The position of the autoinhibitory coiled coil domain (autoN) and H2A.W binding domains have been reported previously (Nartey et al. 2023; Osakabe et al. 2021).

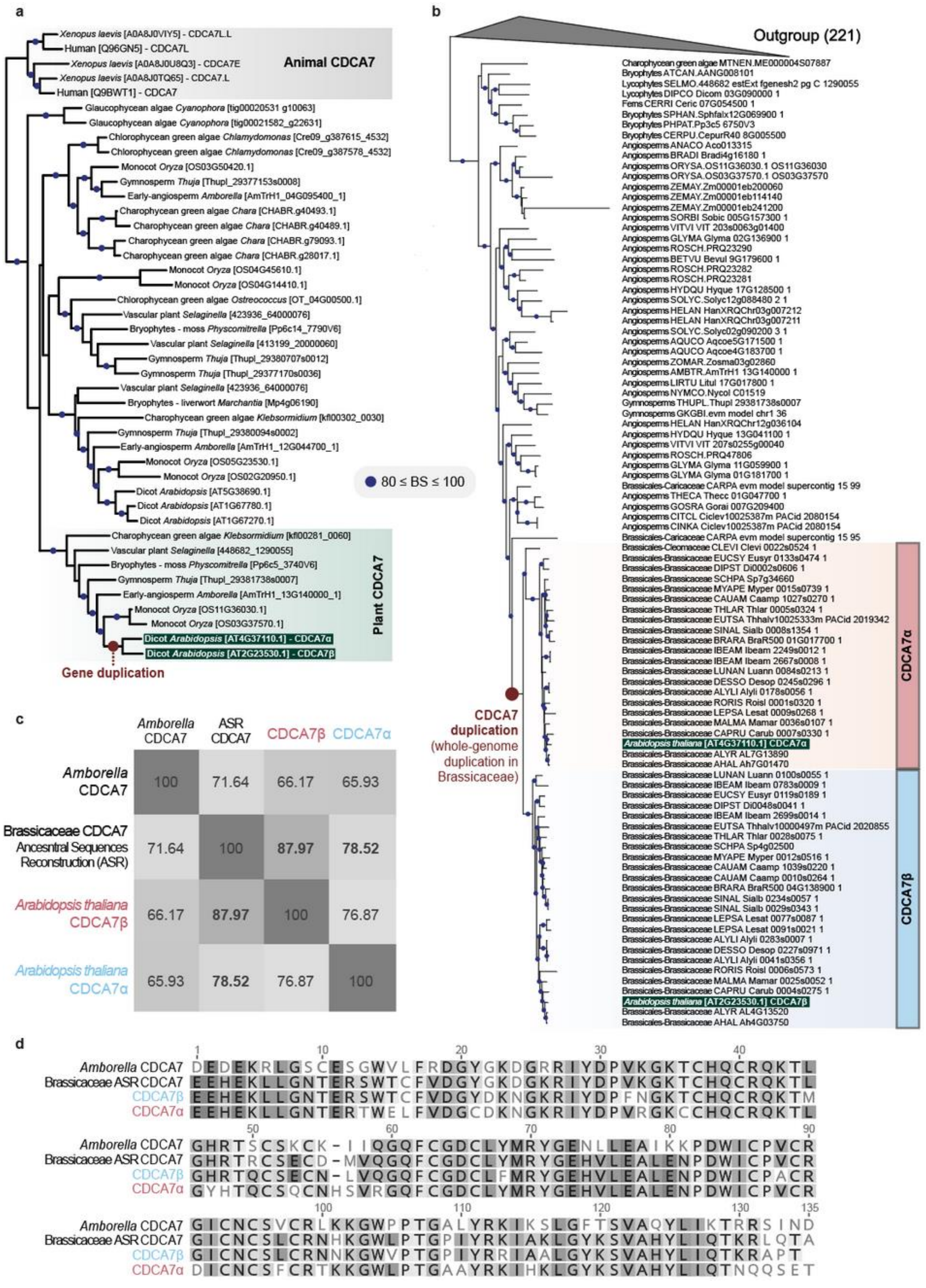

a
Xenopus laevis [A0A8J0VIY5] - CDCA7L.L
Human [Q96GN5] - CDCA7L
Xenopus laevis [A0A8J0U8Q3] - CDCA7E
Xenopus laevis [A0A8J0TQ65] - CDCA7.L
Human [Q9BWT1] - CDCA7
Animal CDCA7
Glaucophycean algae Cyanophora [tig00020531 g10063]
Glaucophycean algae Cyanophora [tig00021582_g22631]
Chlorophycean green algae Chlamydomonas [Cre09_g387615_4532]
Chlorophycean green algae Chlamydomonas [Cre09_g387578_4532]
Monocot Oryza [OS03G50420.1]
Gymnosperm Thuja [Thupl_29377153s0008]
Early-angiosperm Amborella [AmTrH1_04G095400_1]
Charophycean green algae Chara [CHABR.g40493.1]
Charophycean green algae Chara [CHABR.g40489.1]
Charophycean green algae Chara [CHABR.g79093.1]
Charophycean green algae Chara [CHABR.g28017.1]
Monocot Oryza [OS04G45610.1]
Monocot Oryza [OS04G14410.1]
Chlorophycean green algae Ostreococcus [OT_04G00500.1]
Vascular plant Selaginella [423936_64000076]
Bryophytes - moss Physcomitrella [Pp6c14_7790V6]
Vascular plant Selaginella [413199_20000060]
Gymnosperm Thuja [Thupl_29380707s0012]
Gymnosperm Thuja [Thupl_29377170s0036]
Vascular plant Selaginella [423936_64000076]
Bryophytes - liverwort Marchantia [Mp4g06190]
Charophycean green algae Klebsormidium [kfl00302_0030]
Gymnosperm Thuja [Thupl_29380094s0002]
Early-angiosperm Amborella [AmTrH1_12G044700_1]
Monocot Oryza [OS05G23530.1]
Monocot Oryza [OS02G20950.1]
Dicot Arabidopsis [AT5G38690.1]
Dicot Arabidopsis [AT1G67780.1]
Dicot Arabidopsis [AT1G67270.1]
80 ≤ BS ≤ 100
Charophycean green algae Klebsormidium [kfl00281_0060]
Vascular plant Selaginella [448682_1290055]
Bryophytes - moss Physcomitrella [Pp6c5_3740V6]
Gymnosperm Thuja [Thupl_29381738s0007]
Early-angiosperm Amborella [AmTrH1_13G140000_1]
Monocot Oryza [OS11G36030.1]
Monocot Oryza [OS03G37570.1]
Dicot Arabidopsis [AT4G37110.1] - CDCA7α
Dicot Arabidopsis [AT2G23530.1] - CDCA7β
Plant CDCA7
Gene duplication
b
Outgroup (221)
Charophycean green algae MTNEN.ME000004S07887
Bryophytes ATCAN.AANG008101
Lycophytes SELMO.448682 estExt fgenesh2 pg C 1290055
Lycophytes DIPCO Dicom 03G090000 1
Ferns CERRI Ceric 07G054500 1
Bryophytes SPHAN.Sphfalx12G069900 1
Bryophytes PHPAT.Pp3c5 6750V3
Bryophytes CERPU.CepurR40 8G005500
Angiosperms ANACO Aco013315
Angiosperms BRADI Bradi4g16180 1
Angiosperms ORYSA.OS11G36030.1 OS11G36030
Angiosperms ORYSA.OS03G37570.1 OS03G37570
Angiosperms ZEMAY.Zm00001eb200060
Angiosperms ZEMAY.Zm00001eb114140
Angiosperms ZEMAY.Zm00001eb241200
Angiosperms SORBI Sobic 005G157300 1
Angiosperms VITVI VIT 203s0063g01400
Angiosperms GLYMA Glyma 02G136900 1
Angiosperms ROSCH.PRQ23290
Angiosperms BETVU Bevul 9G179600 1
Angiosperms ROSCH.PRQ23282
Angiosperms ROSCH.PRQ23281
Angiosperms HYDQU Hyque 17G128500 1
Angiosperms SOLYC.Solyc12g088480 2 1
Angiosperms HELAN HanXRQChr03g007212
Angiosperms HELAN HanXRQChr03g007211
Angiosperms SOLYC.Solyc02g090200 3 1
Angiosperms AQUCO Aqcoe5G171500 1
Angiosperms AQUCO Aqcoe4G183700 1
Angiosperms ZOMAR.Zosma03g02860
Angiosperms AMBTR.AmTrH1 13G140000 1
Angiosperms LIRTU Litul 17G017800 1
Angiosperms NYMCO.Nycol C01519
Gymnosperms THUPL.Thupl 29381738s0007
Gymnosperms GKGBI.evm model chr1 36
Angiosperms HELAN HanXRQChr12g036104
Angiosperms HYDQU Hyque 13G041100 1
Angiosperms VITVI VIT 207s0255g00040
Angiosperms ROSCH.PRQ47806
Angiosperms GLYMA Glyma 11G059900 1
Angiosperms GLYMA Glyma 01G181700 1
Brassicales-Caricaceae CARPA evm model supercontig 15 99
Angiosperms THECA Thecc 01G047700 1
Angiosperms GOSRA Gorai 007G209400
Angiosperms CITCL Ciclev10025387m PACid 2080154
Angiosperms CINKA Ciclev10025387m PACid 2080154
Brassicales-Caricaceae CARPA evm model supercontig 15 95
Brassicales-Cleomaceae CLEVI Clevi 0022s0524 1
Brassicales-Brassicaceae EUCSY Eusyr 0133s0474 1
Brassicales-Brassicaceae DIPST Di0002s0606 1
Brassicales-Brassicaceae SCHPA Sp7g34660
Brassicales-Brassicaceae MYAPE Myper 0015s0739 1
Brassicales-Brassicaceae CAUAM Caamp 1027s0270 1
Brassicales-Brassicaceae THLAR Thlar 0005s0324 1
Brassicales-Brassicaceae EUTSA Thhalv10025333m PACid 2019342
Brassicales-Brassicaceae SINAL Sialb 0008s1354 1
Brassicales-Brassicaceae BRARA BraR500 01G017700 1
Brassicales-Brassicaceae IBEAM Ibeam 2249s0012 1
Brassicales-Brassicaceae IBEAM Ibeam 2667s0008 1
Brassicales-Brassicaceae LUNAN Luann 0084s0213 1
Brassicales-Brassicaceae DESSO Desop 0245s0296 1
Brassicales-Brassicaceae ALYLI Alyli 0178s0056 1
Brassicales-Brassicaceae RORIS Roisl 0001s0320 1
Brassicales-Brassicaceae LEPSA Lesat 0009s0268 1
Brassicales-Brassicaceae MALMA Mamar 0036s0107 1
Brassicales-Brassicaceae CAPRU Carub 0007s0330 1
Arabidopsis thaliana [AT4G37110.1] CDCA7α
Brassicales-Brassicaceae ALYR AL7G13890
Brassicales-Brassicaceae AHAL Ah7G01470
Brassicales-Brassicaceae LUNAN Luann 0100s0055 1
Brassicales-Brassicaceae IBEAM Ibeam 0783s0009 1
Brassicales-Brassicaceae EUCSY Eusyr 0119s0189 1
Brassicales-Brassicaceae DIPST Di0048s0041 1
Brassicales-Brassicaceae IBEAM Ibeam 2699s0014 1
Brassicales-Brassicaceae EUTSA Thhalv10000497m PACid 2020855
Brassicales-Brassicaceae THLAR Thlar 0028s0075 1
Brassicales-Brassicaceae SCHPA Sp4g02500
Brassicales-Brassicaceae MYAPE Myper 0012s0516 1
Brassicales-Brassicaceae CAUAM Caamp 1039s0220 1
Brassicales-Brassicaceae CAUAM Caamp 0010s0264 1
Brassicales-Brassicaceae BRARA BraR500 04G138900 1
Brassicales-Brassicaceae SINAL Sialb 0234s0057 1
Brassicales-Brassicaceae SINAL Sialb 0029s0343 1
Brassicales-Brassicaceae LEPSA Lesat 0077s0087 1
Brassicales-Brassicaceae LEPSA Lesat 0091s0021 1
Brassicales-Brassicaceae ALYLI Alyli 0283s0007 1
Brassicales-Brassicaceae DESSO Desop 0227s0971 1
Brassicales-Brassicaceae ALYLI Alyli 0041s0356 1
Brassicales-Brassicaceae RORIS Roisl 0006s0573 1
Brassicales-Brassicaceae MALMA Mamar 0025s0052 1
Brassicales-Brassicaceae CAPRU Carub 0004s0275 1
Arabidopsis thaliana [AT2G23530.1] CDCA7β
Brassicales-Brassicaceae ALYR AL4G13520
Brassicales-Brassicaceae AHAL Ah4G03750
CDCA7α
CDCA7β
CDCA7 duplication (whole-genome duplication in Brassicaceae)
c
Amborella CDCA7
ASR CDCA7
CDCA7β
CDCA7α
Amborella CDCA7
100
71.64
66.17
65.93
Brassicaceae CDCA7 Ancesntral Sequences Reconstruction (ASR)
71.64
100
87.97
78.52
Arabidopsis thaliana CDCA7β
66.17
87.97
100
76.87
Arabidopsis thaliana CDCA7α
65.93
78.52
76.87
100
d
1
10
20
30
40
Amborella CDCA7 DEDEKRLGSCESGWVLFRDGYGKDGRRIYDPVKGKTCHQCRQKTL
Brassicaceae ASR CDCA7 EEHEKLLGNTERSWTCFVDGYGKDGKRIYDPVKGKTCHQCRQKTL
CDCA7β EEHEKLLGNTERSWTCFVDGYDKNGKRIYDPFNGKTCHQCRQKTM
CDCA7α EEHEKLLGNTERTWELFVDGCDKNGKRIYDPVRGKCCHQCRQKTL
50
60
70
80
90
Amborella CDCA7 GHRTSCSKCK-IIQGQFCGDCLYMRYGENLLEAIKKPDWICPVCR
Brassicaceae ASR CDCA7 GHRTRCSECD-MVQGQFCGDCLYMRYGEHVLEALENPDWICPVCR
CDCA7β GHRTQCSECN-LVQGQFCGDCLFMRYGEHVLEALENPDWICPACR
CDCA7α GYHTQCSQCNHSVRGQFCGDCLYMRYGEHVLEALENPDWICPVCR
100
110
120
130
135
Amborella CDCA7 GICNCSVCRLKKGWPPTGALYRKIKSLGFTSVAQYLIKTRRSIND
Brassicaceae ASR CDCA7 GICNCSLCRNHKGWLPTGPIYRKIAKLGYKSVAHYLIQTKRLQTA
CDCA7β GICNCSLCRNNKGWVPTGPIYRRIAALGYKSVAHYLIQTKRAPT
CDCA7α DICNCSFCRTKKGWLPTGAAYRKIHKLGYKSVAHYLIQTNQQSET

**Figure S3. The evolutionary relationships of CDCA7 proteins across different evolutionary scales.**

**a**, The phylogeny of CDCA7 proteins among representative taxa of Archaeplastida. Xenopus and human CDCA7 proteins provided as an outgroup, previously identified as homologs of plant CDCA7 (Funabiki et al, 2023). **b**, The phylogeny of CDCA7 proteins among representative taxa of Viridiplantae, including multiple species from Brassicaceae. The phylogeny indicates that A. thaliana CDCA7α [AT4G37110.1] and A. thaliana CDCA7β [AT2G23530.1] underwent duplication in the early Brassicaceae. **c**, Protein identity comparison of the zinc finger domains of four CDCA proteins: Arabidopsis CDCA7α & CDCA7β, ancestral sequence reconstruction (ASR) of CDCA7 protein sequence at the node of Brassicaceae CDCA7 duplication (n=48), and *Amborella* CDCA7 as an outgroup. The average posterior probability of ASR sequence (total 406 residues) is 0.916 (n=406, median=0.996, σ=0.151). The zinc finger domain of CDCA7β shows a higher similarity to the ASR sequence compared to CDCA7α, indicating that CDCA7β may be more ancestral than CDCA7α. **d**, ASR of CDCA7 zinc finger aligned with three CDCA7 proteins.

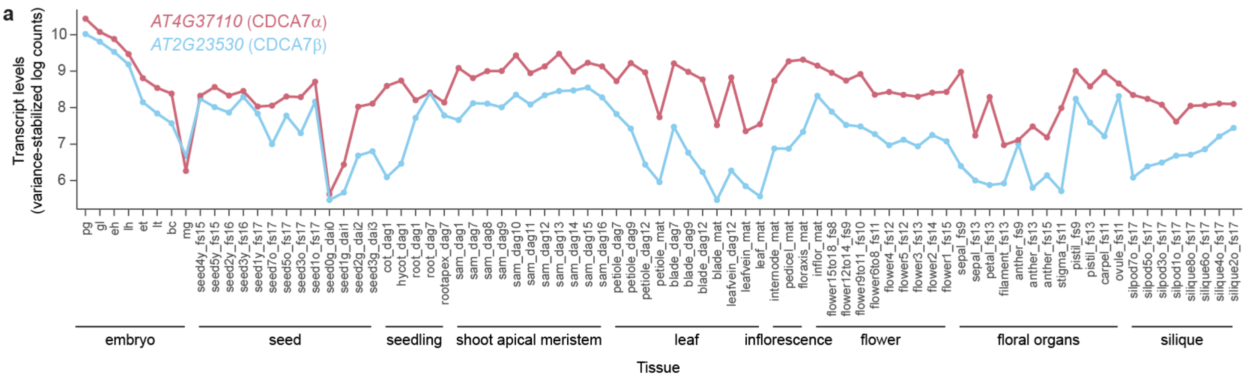


**Figure S4**. **CDCA7 expression in Arabidopsis tissues**

Transcript levels of *AT4G37110* and *AT2G23530* in transcriptome datasets from various developmental stages, shown as variance-stabilized log counts, from Hoffman et al (2019 Plant Reproduction).

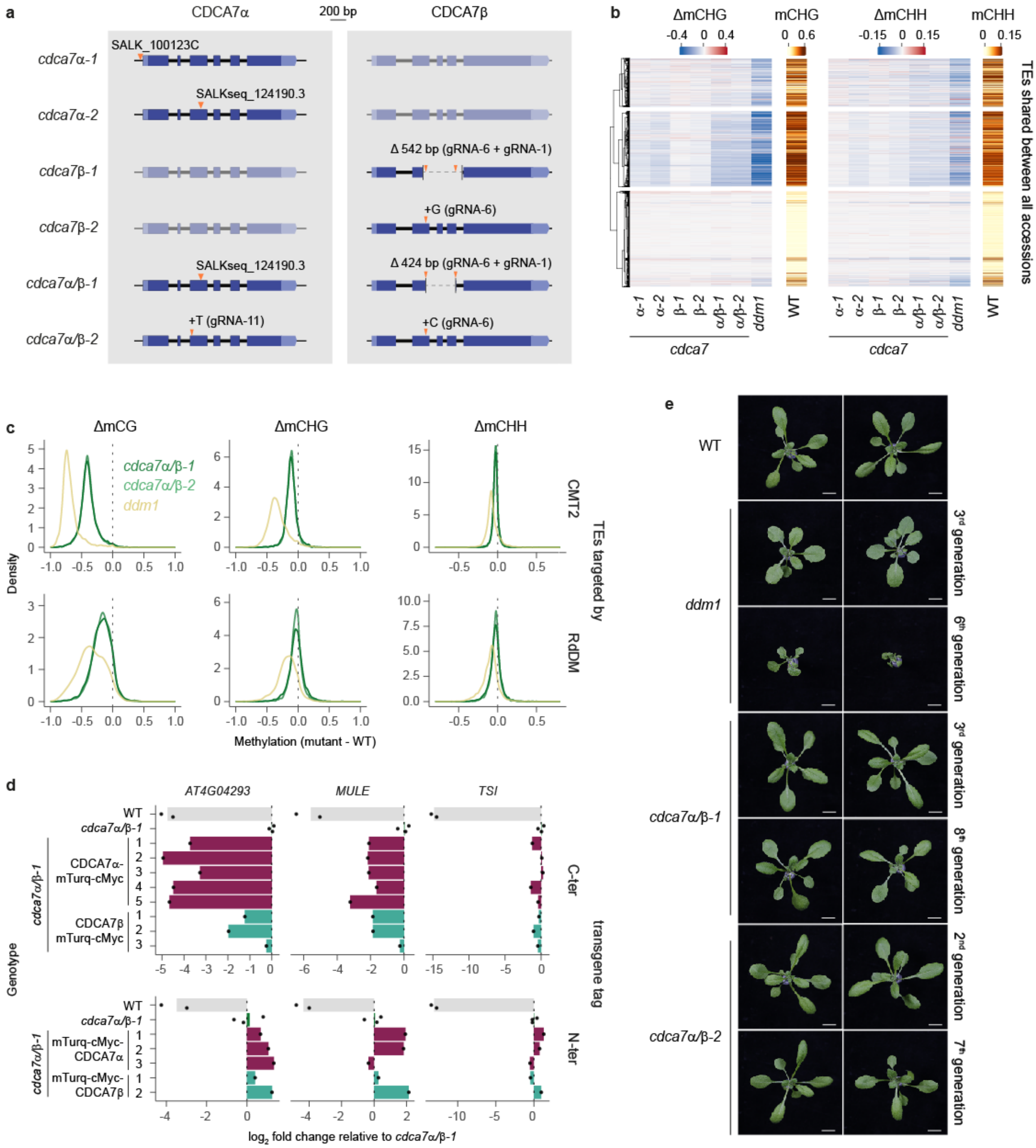


**Figure S5. CDCA7a/b primarily affect the DNA methylation of CMT2-targeted TEs**

**a**, Examples of mutations used in this study, drawn to scale. Additional *cdca7* mutations are listed in Supplementary Table 3. **b,** Heatmap comparing Δm5C (mutant - WT) for *cdca7* null mutations. K-mean clusters were defined based on ΔmCG (**Fig. 2c**). The WT value is shown as a reference. **c**, Distributions showing 5mC differences between the indicated mutants and the WT in CG, CHG and CHH contexts, distinguishing TEs targeted by CMT2 or RdDM. **d**, RT-qPCR quantification of transcripts from heterochromatic loci in cdca7α/β mutants complemented with transgene containing either a C or N-terminal tag. For each transgene, primary transformants were plotted individually to account for differences in transgene expression. Barplots show means, with black points showing biological replicates. Expression levels are normalized to cdca7α/β-1. **e**, Developmental phenotypes of the indicated mutants after 20 days of growth in long days. Scale bar : 10 mm.

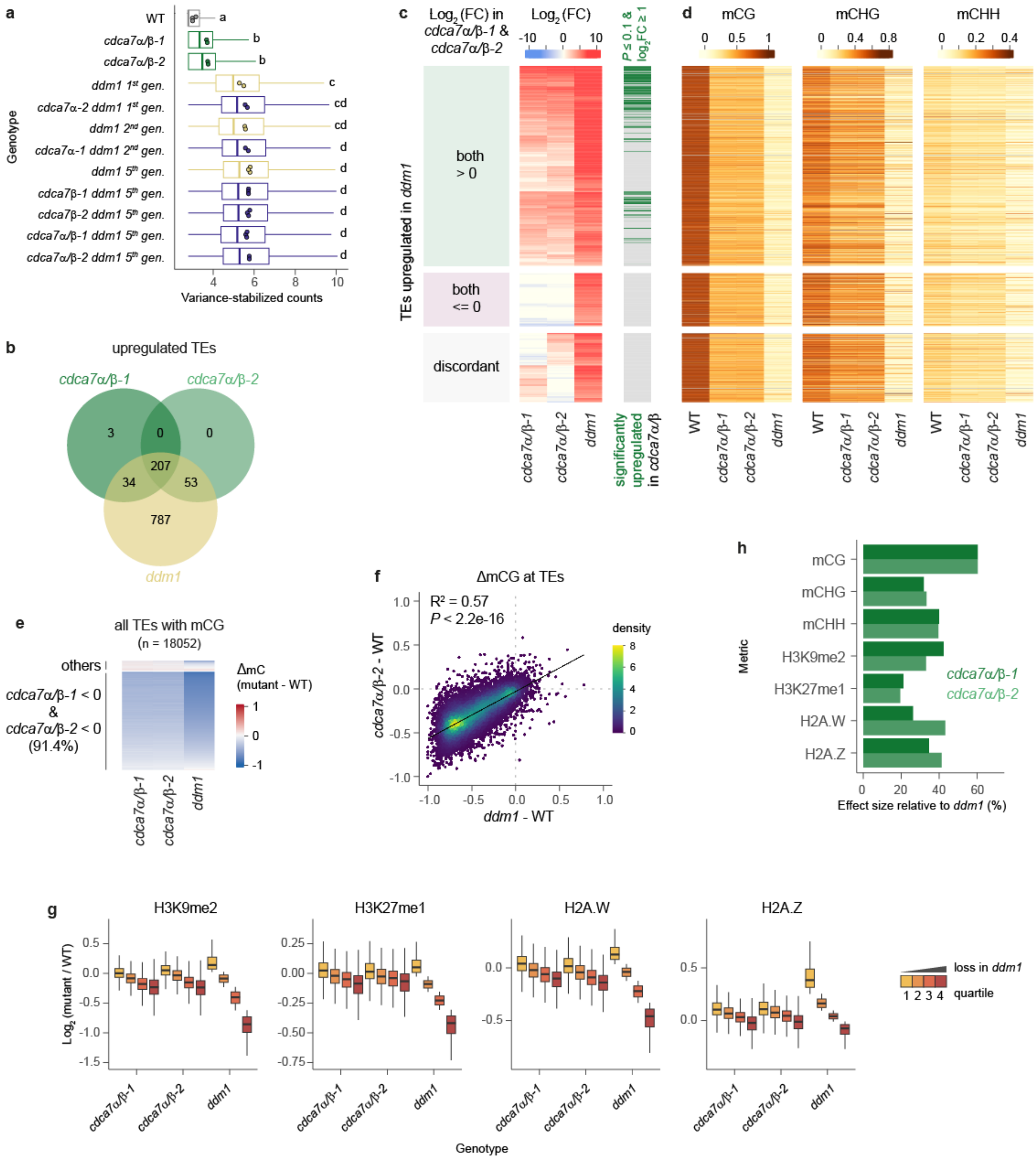


**Figure S6. CDCA7α and CDCA7β function through DDM1 to maintain mCG.**

**a**, Variance-stabilized transcript levels of transposable elements (TEs) upregulated in any of the *ddm1* mutants (yellow, n = 1147). Mean values of biological replicates are shown as circles, with statistical groups determined by Tukey's HSD test ($P < 0.05$) denoted by lowercase letters. **b**, Overlap of upregulated TEs between mutants. **c**, **d**, Effects of *cdca7α/β* mutations on transcript levels (**c**) and 5mC (**d**) at DDM1-silenced TEs. TEs are grouped by $\log_2$ fold change ($\log_2$(FC)) in *cdca7α/β-1* and *cdca7α/β-2* (left), with those meeting upregulation thresholds in both mutants highlighted in green. **e**, mCG changes, using all TEs with initial mCG > 5%. **f**, Correlation of mCG loss at TEs in *cdca7α/β-2* relative to *ddm1*. Pearson correlation coefficients ($R^2$) and a permutation test-derived *P*-value are shown. **g,** Changes in histone marks and variants at TEs, stratified into quartiles based on the change observed in *ddm1*. **h**, Effect size of *cdca7α/β* mutations relative to *ddm1* across various metrics.

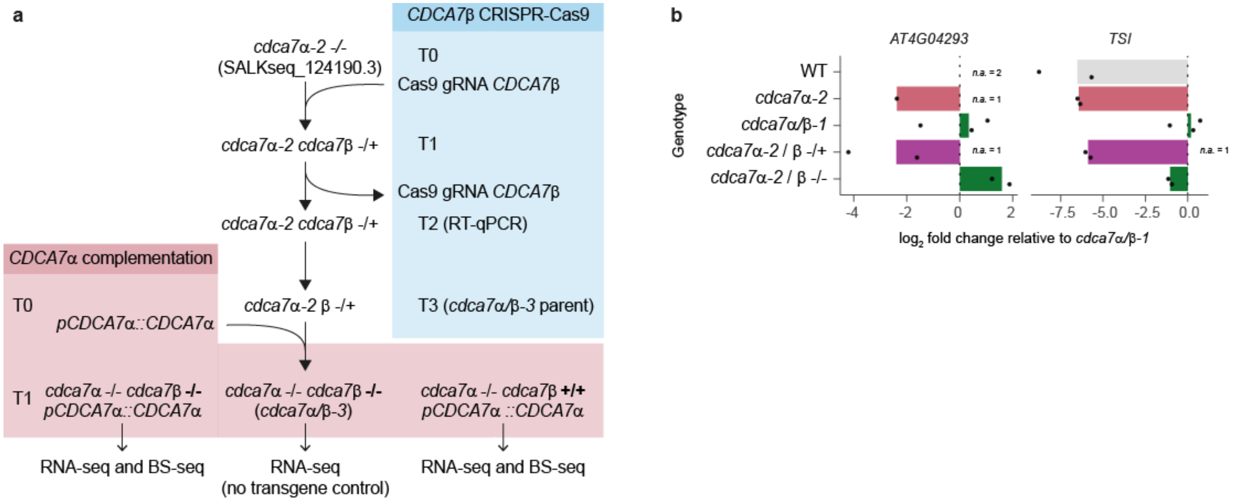


**Figure S7. Complementation strategy of *cdca7α/β* mutants.**

**a**, Illustration of the genetic scheme used to generate and complement cdca7α/β sesquimutants. We mutagenized *CDCA7β* in the *cdca7α-2* mutant background with CRISPR-Cas9. After isolating *cdca7β* mutations in T1, we selected T2 lines with no CRISPR-Cas9 transgene and verified that one *CDCA7β* allele was sufficient to maintain silencing (shown in **b**.). T3 sesquimutants were complemented with a pCDCA7α::CDCA7α-mTurq-cMyc transgene. T1 transformants were isolated with *CDCA7β* homozygous mutant or WT, and we generated transcriptomes (**Fig. 4, 5**) and methylomes (**Fig. 5**). **b**, RT-qPCR quantification of two TE transcripts in *cdca7α/β* mutants, represented as in **a**. In some samples, transcript levels were below the detection limit, indicated by no amplification (n.a.).

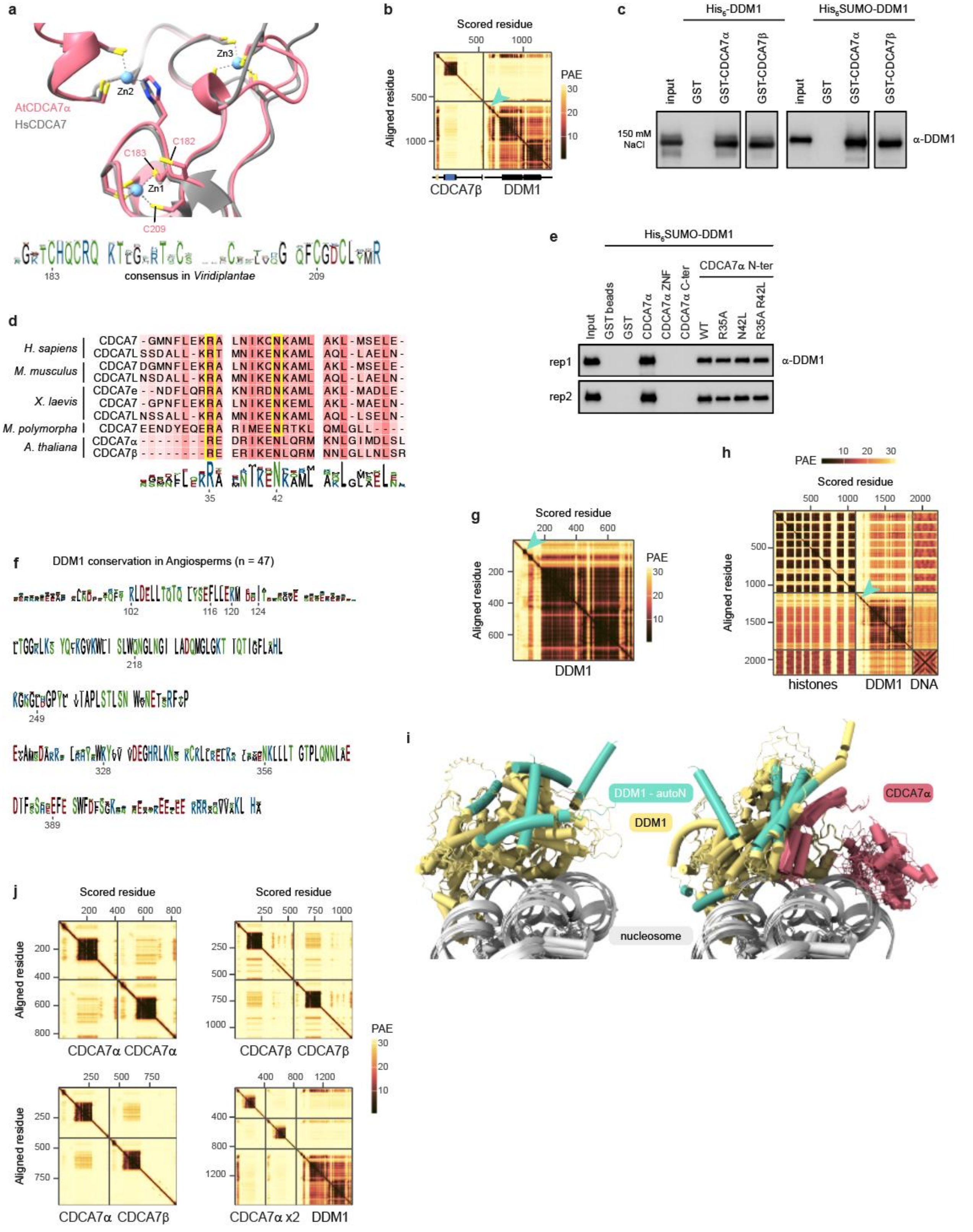


**Figure S8. Physical interaction between CDCA7α/β and DDM1**

**a**, Alphafold3 prediction of *A. thaliana* CDCA7α zinc finger domain aligned to *H. sapiens* CDCA7 crystal structure (PDB ID 8TLK). The zinc atoms and their binding residues are shown. The consensus sequence in Viridiplantae is shown at the bottom (n = 37). **b**, Predicted Aligned Error (PAE) from AlphaFold3, modeling the interaction between CDCA7β and DDM1. Protein domains are drawn at the bottom. A teal arrow shows DDM1 autoN domain. **c**, In vitro co-immunoprecipitation of recombinant CDCA7α and CDCA7β with DDM1, analyzed by Western Blot, at 150 mM NaCl. **d**, Alignment of CDCA7 proteins from model species. A yellow box highlights CDCA7 residues predicted to interact with HELLS/DDM1 from the corresponding species. The position of CDCA7α R35 and N42 is indicated on the consensus sequence. **e**, In vitro co-immunoprecipitation of recombinant DDM1 with mutant truncated CDCA7α. **f**, Conservation in Angiosperms (n = 47) of DDM1 residues predicted to interact with CDCA7α. Numbers indicate residues with predicted contact to CDCA7α. **g**, **h**, PAE plot of DDM1 alone (**g**) or with a nucleosome (**h**). A teal arrow shows DDM1 autoN domain. **i**, Prediction of the DDM1-nucleosome structure with (right) or without (left) CDCA7α, showing inconsistency between the five superimposed Alphafold3 models.

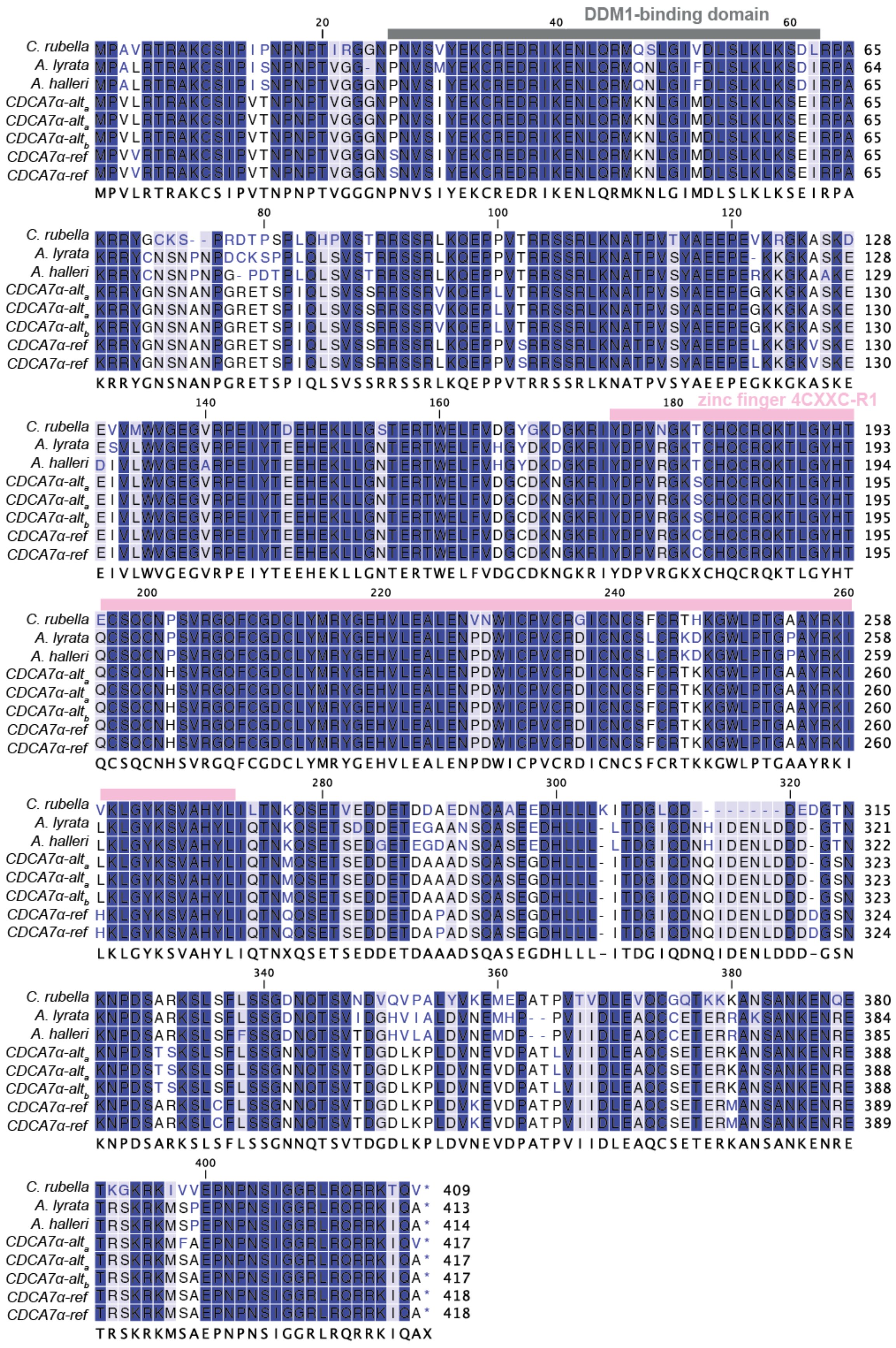
DDM1-binding domain
zinc finger 4CXXC-R1
C. rubella
A. lyrata
A. halleri
CDCA7α-alt
CDCA7α-ref
MPVLRTRAKCSIPVTNPNPTVGGGNPNVSIYEKCREDRIKENLQRMKNLGIMDLSLKLKSEIRPA
KRRYGNSNANPGRETSPIQLSVSSRRSSRLKQEPPVTRRSSRLKNATPVSYAEEPEGKKGKASKE
EIVLWVGEGVRPEIYTEEHEKLLGNTERTWELFVDGCDKNGKRIYDPVRGKXCHQCRQKTLGYHT
QCSQCNHSVRGQFCGDCLYMRYGEHVLEALENPDWICPVCRDICNCSFCRTKKGWLPTGAAYRKI
LKLGYKSVAHYLIQTNXQSETSEDDETDAAADSQASEGDHLLL-ITDGIQDNQIDENLDDD-GSN
KNPDSARKSLSFLSSGNNQTSVTDGDLKPLDVNEVDPATPVIIDLEAQCSETERKANSANKENRE
TRSKRKMSAEPNPNSIGGRLRQRRKIQAX

**Figure S9. Protein sequences of *CDCA7α* from *A. thaliana* haplotypes and other Brassicaceae**
Exon regions were extracted from three *CDCA7*α alleles and relative species based on a gene model of AtRTD3. The amino acids under the alignment represent consensus sequences. Rectangles on the consensus indicate the segregating sites between alleles within the conserved domain. The sequences of *CDCA7*α-alt$_a$ (chr4:17486863$_{\text{non-ref}}$) are extracted from 9888 and 6069, the sequence of *CDCA7α-alt$_b$* (chr4:17497441$_{\text{non-ref}}$) is from 9543, and the sequences of *CDCA7*α-ref are extracted from 6909 (Col-0) and 8236.

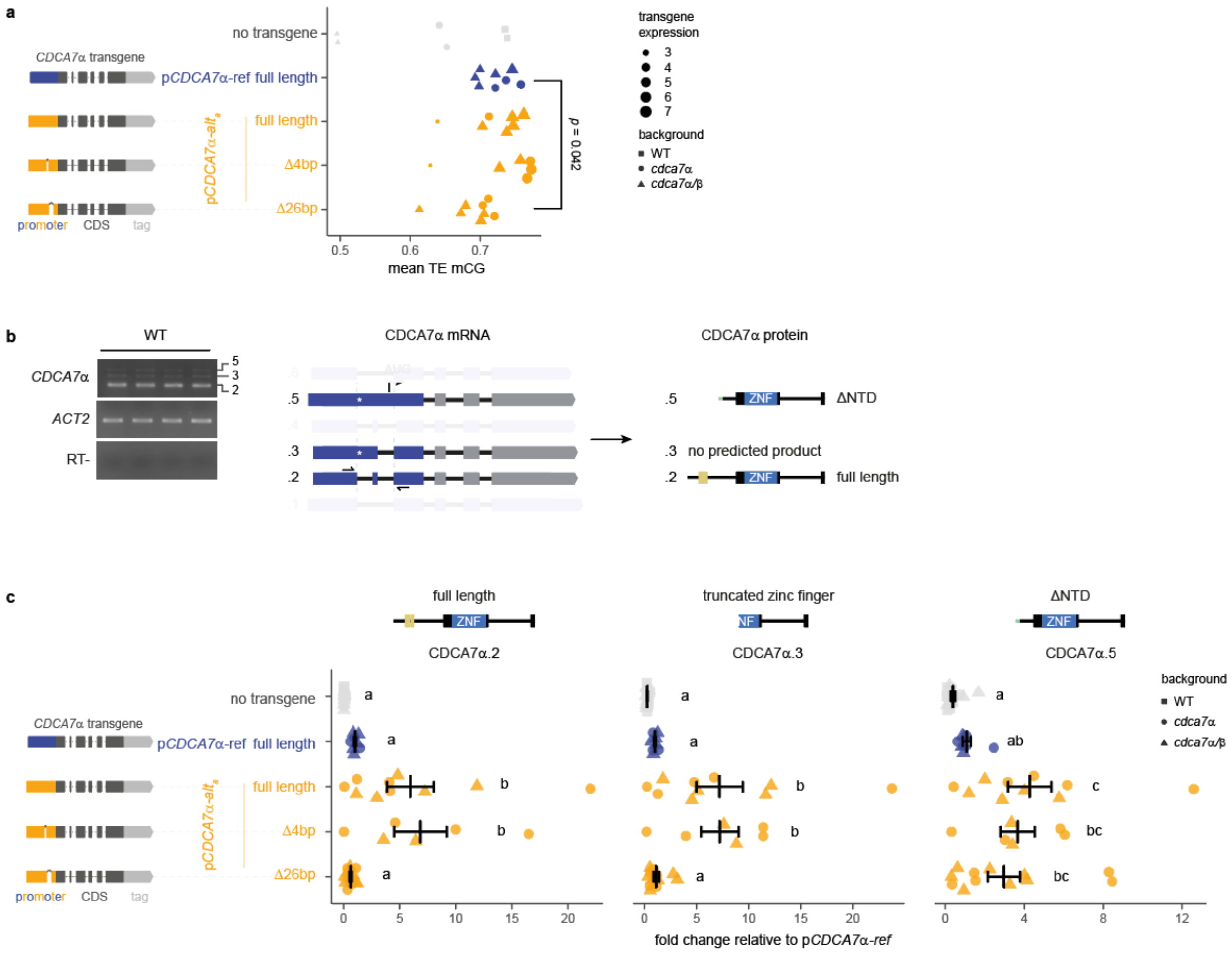


**Figure S10. The 26 bp indel in the *CDCA7α* promoter regulates transcript isoform usage**
**a**, Mean mCG at TEs in complemented *cdca7α/β* and *cdca7α* mutants. Each data point is an individual plant, with point size indicating transgene expression levels. The *p-value* is from a two-sided Student's test. **b**, CDCA7α transcripts isoforms detected by RT-PCR in four biological replicates. The *ACTIN2* gene was amplified as a loading control, and a negative control with no reverse transcription is shown (RT-). Matching annotated isoforms ((R. Zhang et al. 2022), confirmed by Sanger sequencing, are highlighted (middle). Primer positions are indicated by arrows, and translation start (AUG) and stop (*) codons are shown. Corresponding protein products are shown (right). **c**, Quantification of *CDCA7*α transcripts isoforms by RT-qPCR. Each data point is an individual primary transformant with mean

values ± standard error shown in black and significance between groups indicated by lowercase letters (Tukey's HSD tests, $p < 0.05$). Expression levels are normalized to mutants complemented with pCDCA7α-ref.

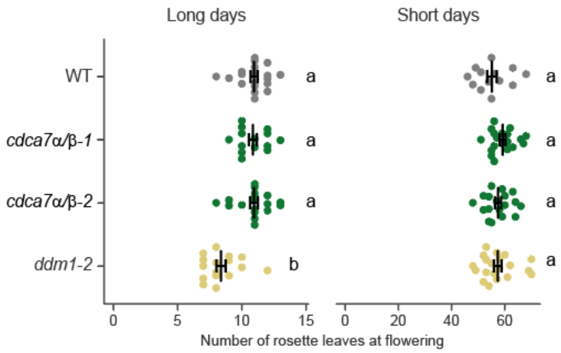


**Figure S11. Flowering time of *cdca7α/ß* null mutants.**
Flowering time, measured as developmental stage at flowering (n = 12 to 19). The statistical groups, indicated by lowercase letters, were determined by one-way ANOVA with Tukey-Kramer post-hoc tests ($P < 1e\text{-}05$).
